# Astrocytic Regulation of Basal Ganglia Dopamine/D2-Dependent Behaviors

**DOI:** 10.1101/2021.05.11.443394

**Authors:** Rosa Mastrogiacomo, Gabriella Trigilio, Daniel Dautan, Céline Devroye, Valentina Ferretti, Enrica Vitali, Genny Orso, Roberto Marotta, Federica Maltese, Gessica Piras, Alessia Forgiarini, Giada Pacinelli, Debora A. Rothmond, John L. Waddington, Filippo Drago, Maria Antonietta De Luca, Gian Marco Leggio, Cynthia S. Weickert, Francesca Managò, Francesco Papaleo

**Affiliations:** Genetics of Cognition laboratory, Neuroscience area, Istituto Italiano di Tecnologia, via Morego, 30, 16163 Genova, Italy; Department of Biomedical and Biotechnological Sciences, University of Catania, Catania, Italy; Fondazione IRCCS Ca’ Granda Ospedale Maggiore Policlinico, Milano, Italy; Department of Pharmaceutical and Pharmacological Sciences, University of Padova, Padova, Italy; Department of Biomedical Sciences, University of Cagliari, Cagliari, Italy; Schizophrenia Research Laboratory, Neuroscience Research Australia, Sydney, Australia; School of Pharmacy and Biomolecular Sciences, Royal College of Surgeons in Ireland, Dublin 2, Ireland

**Keywords:** Astrocytes, dopamine, dysbindin-1A, D2 receptor, schizophrenia, globus pallidus external segment, motivation, sensorimotor gating, substantia nigra pars compacta

## Abstract

Astrocytic involvement in dopamine dynamics related to motivational and sensorimotor gating abilities is unknown. We found that dysbindin-1 (Dys1) hypofunction increases the activity of astrocytes, which express only the isoform Dys1A that is reduced in the caudate of patients with schizophrenia. Astrocytic Dys1A disruption resulted in avolition and sensorimotor gating deficits, increased astrocytic dopamine D2 receptors and decreased dopaminergic tone within basal ganglia. Notably, astrocytic Dys1A hypofunction disrupted dopamine dynamics linked to reward expectancy and interconnected with astrocytes Ca^2+^ responses mainly in the globus pallidus externus (GPe). Finally, we proved these phenotypes were mediated by D2 receptors in astrocytes as their selective deletion in astrocytes either in GPe or SNc/VTA enhanced motivation and sensorimotor gating abilities as well as dopaminergic release in the GPe. Therefore, astrocytes control motivational and sensorimotor gating processes through Dys1A/D2-dependent mechanisms within the basal ganglia.

## Introduction

Astrocytes fundamentally contribute to brain homeostasis and play a role in brain physiology through reciprocal communication with neuronal cells ^1–6^. Neurotransmitters can modulate astrocytic activity and, *vice versa*, astrocytic gliotransmitters can regulate neurotrasmission ^2, 4, 5, 7, 8^. Astrocytic-neuronal communication has been consistently reported to involve glutamatergic, cannabinoid, ATP/adenosine, cholinergic, and GABAergic systems ^2^. Notably, an active role for astrocytes in dopamine signaling has emerged ^3, 5, 9, 10^.

The astrocyte-dopamine link appears to rely on distinct molecular factors in prefrontal cortex (PFC), nucleus accumbens (NAcc), globus pallidus external segment (GPe), substantia nigra (SN), and hippocampus ^3, 5, 9–11^. This presupposes heterogeneous astrocytic modulation of dopaminergic signaling with area-specific functions ^12–14^. However, the machinery underlying astrocytic control of dopaminergic signaling has not been fully explored. Moreover, the *in vivo* link between astrocytic activity, dopamine dynamics, and specific behavioral functions remain unknown.

Here, we report the unexpected discovery that the dysbindin-1A (Dys1A) spliced transcript of the Dystrobrevin Binding Protein 1 (DTNBP1) gene is uniquely implicated in the astrocytic regulation of basal ganglia dopamine/D2 signaling and related behavioral processes. Dys1 exists in at least three spliced transcripts, Dys1A, 1B, and 1C ^15^, with 1A and 1C being orthologues in humans and mice ^15–17^. These isoforms are believed to have distinct functions as they are differentially distributed in brain synaptosomes, are present in different functional domains, and have distinct binding partners ^17–19^. However, the specific contribution of each Dys1 isoform in physiological functions, especially astrocyte activity and related behavioral outcomes was yet unexplored.

We show that clinically-relevant Dys1 genetic variations alter astrocyte activity. Specifically, we find that Dys1A is the only isoform expressed in astrocytes and is preferentially involved in astrocytic, but not neuronal, intracellular trafficking. Notably, selective disruption of Dys1A, induces behavioral and dopaminergic alterations related to basal ganglia. This might be clinically relevant as we found that Dys1A is decreased in the caudate of patients with schizophrenia. Finally, using a combination of *in vivo* genetics, Ca^2+^, and dopamine sensor tools, we demonstrate that selective disruption of Dys1A in astrocytes is sufficient to generate distinct motivational, sensorimotor gating, and basal ganglia dopaminergic alterations related to astrocytic D2 receptor mechanisms within the SNc/VTA to GPe circuit. Overall, we show a hitherto unknown mechanism of astrocyte-dopamine interaction in the basal ganglia mediating motivational and sensorimotor gating abilities relevant to schizophrenia and other illnesses that impact motivation and behavioral responses to salient stimuli.

## Results

### Dys1 hypofunction alters astrocytic activity

Dys1 has been linked to neuronal dopaminergic and glutamatergic signaling ^17, 20–24^. However, drosophila dysbindin (dDys) has shown dichotomic regulation of glutamatergic and dopaminergic transmission, with the latter involving glial cells ^25^. To investigate if Dys1 regulates astrocytic activity in mammals, we used Dys1 heterozygous mice (Dys1+/-), a model with direct translational validity for both healthy human subjects and patients with schizophrenia ^20, 26^.

Unbiased microarray analysis showed increased expression of reactive astrogliosis-related genes in Dys1 knockout mice compared to wild-type Dys1+/+ (Fig. 1a and Supplementary Fig. S1). This was confirmed by higher immunoreactivity of the astroglial marker glial fibrillary acid protein (GFAP) in Dys1+/- compared to Dys1+/+ littermates (Fig. 1b), which was similarly evident in PFC, NAcc, dorsal striatum (STR), and GPe (Fig. 1c).

**Figure 1.**
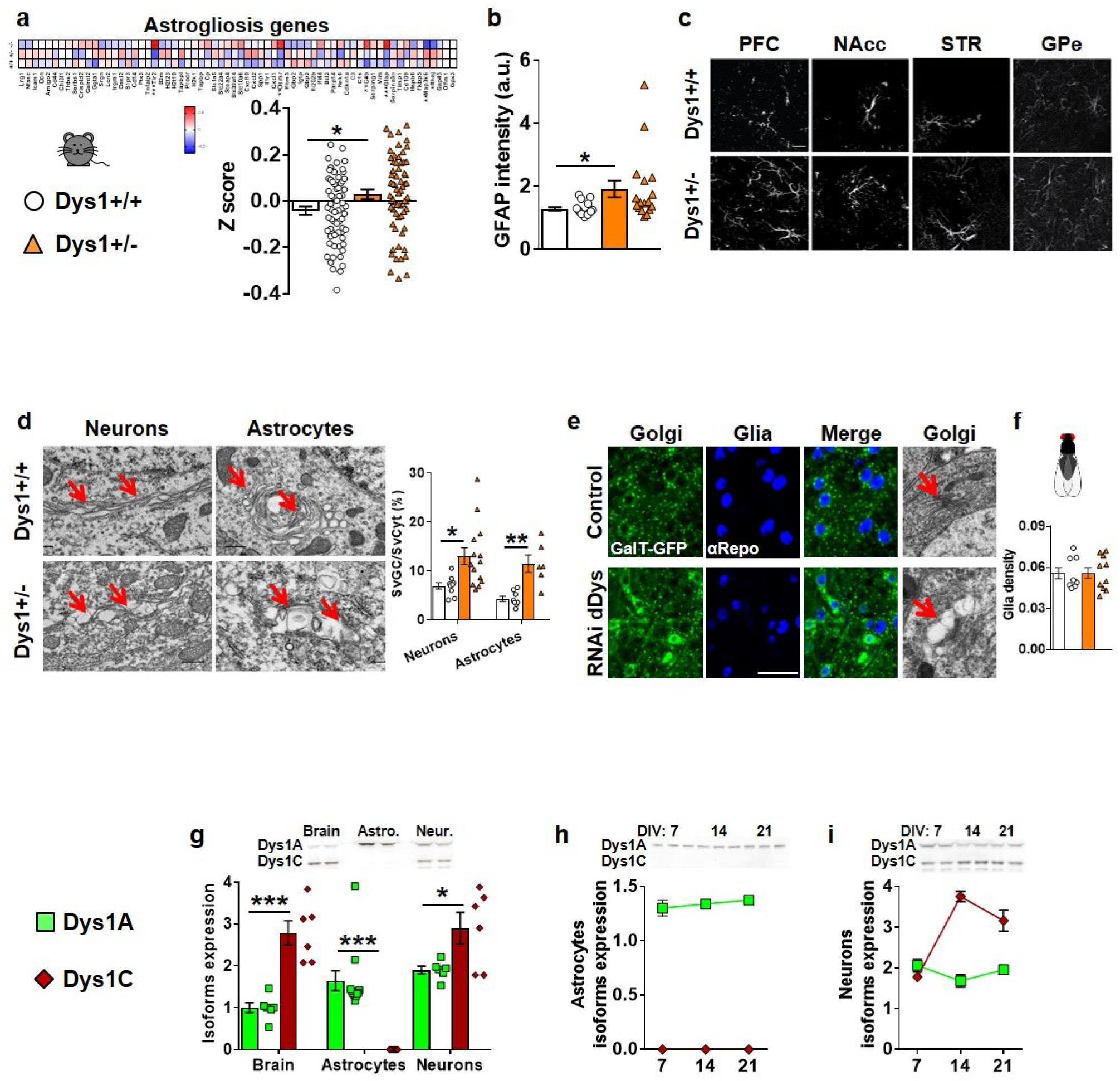
Dys1 is involved in astrocytic activity. **a.** Heat map of 65 inflammatory markers selected by a microarray screening from Dys1+/+ (n24), and Dys1+/- (n25) littermates. The heat map is based on hierarchical clustering of genes involved in inflammation states. All gene expression levels were transformed to scores ranging from −0.5 to 0.5 and were colored blue, white, or red to represent low, moderate, or high expression levels, respectively. The relative expression levels were scaled based on their mean and do not represent expression levels in comparison with controls. Dys1+/- mice show higher expression for these genes compared to Dys1+/+ littermates (t-test: t_128_=-2.23, p=0.028). *p<0.05 vs Dys1+/+. **b.** Quantification of cumulative GFAP intensity from confocal images from PFC, NAcc, STR and GPe displayed by Dys1+/+ and Dys1+/- littermates (n4/genotype, 1/brain region averaged from 9 samples each). Scale bars, 20μm (t-test: t_31_=-2.32, p=0.027) *p<0.05 vs Dys1+/+. **c.** Representative confocal images of GFAP positive astrocytes in the analyzed brain regions. **d.** Transmission electron microscopy (TEM) images of the Golgi Complex (GC) in neurons and astrocytes in Dys1+/+ and Dys1+/- littermates. Surface density of GC (SvGC/SvCyt) in Dys1+/- mice was significantly higher than in Dys1+/+ littermates (neurons t_21_=-2.70, p=0.013; astrocytes t_12_=-4.40, p=0.0009). *p<0.05, **p<0.001 vs Dys1+/+. **e.** Maximum intensity projections of ventral ganglion cells, from Drosophila third instar larvae expressing UAS-GalT-GFP to visualize Golgi cisternae, for controls (tubulin-Gal4/+) and UAS-Dysb RNAi. Tissues were labeled with anti αRepo antibody to visualize glial cells. Scale bar 20 µm. On the right TEM images of third instar larvae brain showing the Golgi apparatus in the neuronal cell bodies of ventral ganglion for the above genotypes. Flies expressing UAS-RNAi Dybs ubiquitously showed swelling of largely inflated Golgi cisternae (arrows). Scale bar 500 nm. **f.** Quantification of the glial nuclei distribution in control (tubulin*-* Gal4*/+*) and RNAi Dysb/tubulin*-*Gal4 flies. **g.** Representative western blots and densitometric analysis of Dys1A (50 kDa) and Dys1C (38 kDa) isoforms. β-actin used as loading control. In brain lysate of adult P90 mice both isoforms were revealed, with higher expression for Dys1C compared to Dys1A (t-test: t_10_=-5.77, p=0.0002). Dys1A was the only isoform expressed in glial cells (t-test: t_20_=-6.32, p*<*0.0001). Similar to brain lysate, neuronal cultures show the expression of both isoforms with relative higher levels of Dys1C (t-test: t_10_=-2.57, p=0.02). **h.** Astrocytes cultures at different developmental time points (day 7=DIV7; day 14= DIV14; day 21= DIV21) confirming no expression of Dys1C in astrocytes. **i.** Neuronal cultures at different developmental time points (DIV7, 14 and 21) showing relative higher expression of Dys1C compared to Dys1A. Bar graphs show mean±s.e.m.

Dys1 plays a crucial role in intracellular vesicular trafficking (Marley & von Zastrow 2010; Ji et al. 2009; Ito et al. 2010; Iizuka et al. 2007). Accordingly, electron microscopy analyses of intracellular vesicles revealed irregularly shaped and swollen cisternae of the Golgi complex with enlarged vesicle-like structures in both neuronal and astrocytic cells of Dys1+/- mice (Fig. 1d and Supplementary Video V1). Equivalent results in altered Golgi complex morphology were obtained by knocking down drosophila dDys either ubiquitously (Fig. 1e) or in glial cells (Supplementary Fig. S1). dDys down-regulation did not affect the number of glial cells (Fig. 1f). This suggests that Dys1 can alter intracellular trafficking from and to Golgi in astrocytes.

Overall, these data provide initial evidence that reduced levels of Dys1 alter astrocytic functioning.

### Distinct neuronal/astrocytes expression and developmental patterns of Dys1 isoforms

We then asked whether Dys1 isoforms might be differentially expressed in neurons and astrocytes. Dys1A was expressed in adult mouse brain, in cultured astrocyte-enriched glial cells, and in cultured neuronal cells (Fig. 1g). In contrast, Dys1C was missing in astrocyte-enriched glial cells, while its expression was higher than Dys1A in the brain as well as in cultured neuronal cells (Fig. 1g). Dys1A showed a stable expression over time in glial and neuronal cultures, while Dys1C was always absent in astrocytes cultures and increased its expression over time in neuronal cells (Fig. 1h,i). We confirmed divergent developmental patterns of expression of these two Dys1 isoforms, with similar findings in human and mouse brains. In particular, samples of human dorsolateral PFC revealed higher Dys1A expression in the embryonic phase, which gradually decreased across development (Supplementary Fig. 1). Conversely, Dys1C expression was lower in embryonic and childhood stages and then increased from adolescence (Supplementary Fig. 1). Similarly, in mouse PFC the expression of Dys1A protein decreased from the embryonic phase, while Dys1C increased its expression in adolescence (Supplementary Fig. 1).

Overall, these data show a similar developmental pattern of Dys1 isoforms expression between mice and humans, and define a previously unexpected constraint of Dys1A expression in astrocytes.

### Dys1A hypofunction alters Golgi complex morphology in astrocytes, but not in neurons

The unique expression of Dys1A isoform in astrocytes prompted us to explore the effects of selective disruption of the Dys1A isoform using a mouse line with flanking LoxP sites targeted to the exon 5 of Dtnbp1 on chromosome 13a (Dys1A^flox/flox^), backcrossed with a germline Cre deleter mouse line ^27^.

Dys1A+/- and Dys1A-/- mice have a gene dosage-dependent reduction and lack of Dys1A isoform, respectively, with unaltered Dys1C expression (Fig. 2a and Supplementary Fig. S2). Notably, in contrast to total Dys1 disruption (Fig. 1d), deletion of only Dys1A disrupted Golgi complex morphology in astrocytic cells, but not in neurons (Fig. 2b).

**Figure 2.**
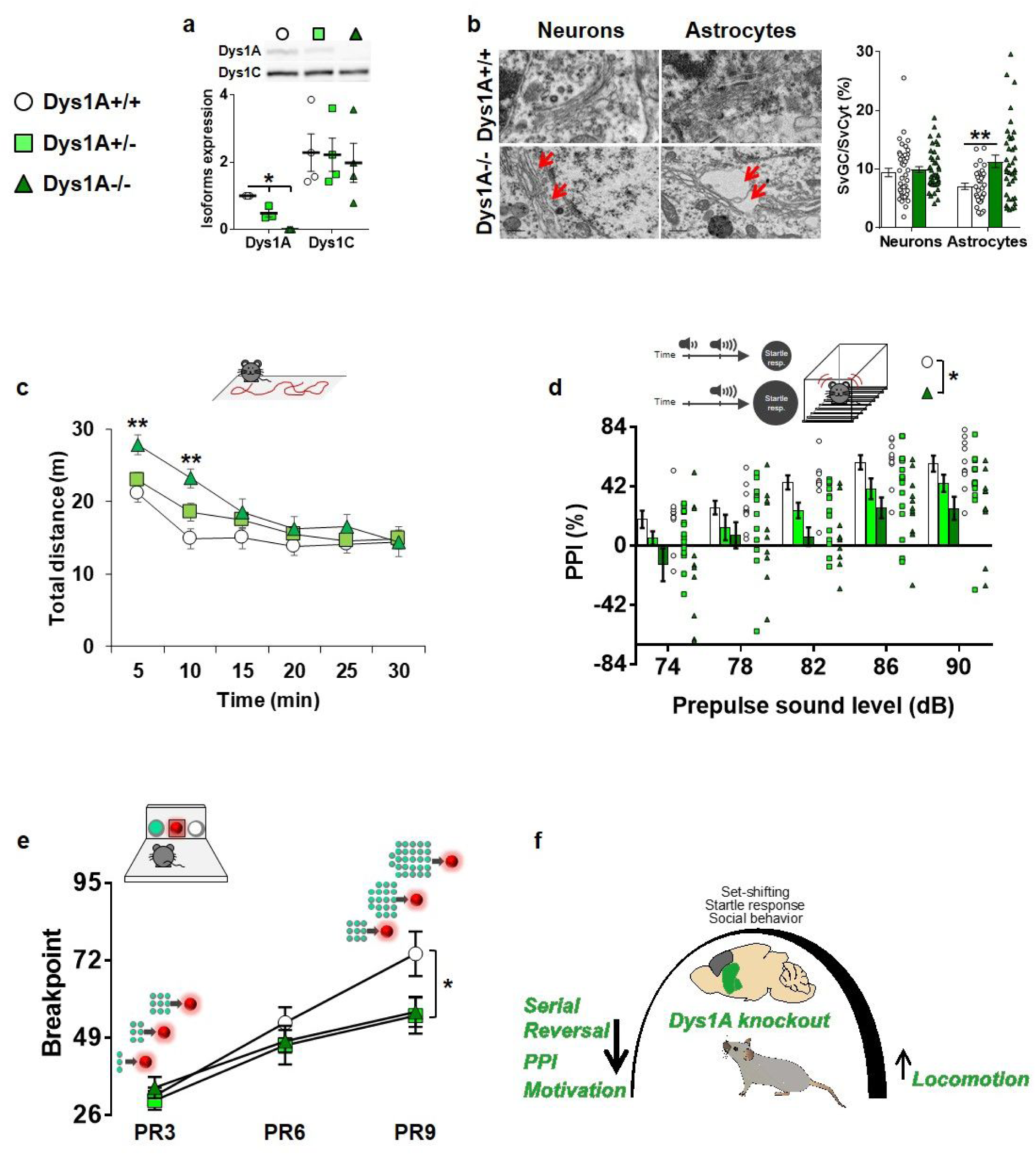
Dys1A disruption impairs basal ganglia-, but not PFC-dependent, behaviors. **a.** Selective reduction of Dys1A does not affect Dys1C expression. Western blots and densitometric analysis in Dys1A+/+, +/- and -/- littermates. Expression of Dys1A (50 kDa), Dys1C (38 kDa). β-actin used as loading control. Expression of Dys1A is reduced in Dys1A+/- and absent in Dys1A-/- in PFC (One-way ANOVA, F_2,6_=60.48; p<0.0005). *p<0.01 vs Dys1A+/+ littermates. Expression of Dys1C was intact across all genotypes (One-way ANOVA, F_2,9_=0.09; p=0.92). n4 mice/group. **b.** Representative transmission electron microscopy (TEM) images of the Golgi Complex (GC, red arrows) from Dys1A+/+ and Dys1A-/- littermates in neuronal and astrocytic cells. Surface density of GC (SvGC/SvCyt) in Dys1A-/- was not altered in neurons (t-test: t_80_=-0.54, p*=*0.59), but significantly increased compared with Dys1A+/+ control mice in astrocytic cells (t-test: t_68_=-3.38, p*<*0.001). **p<0.001 vs Dys1A+/+ littermates. **c.** Spontaneous distance traveled by Dys1A+/+ (n17), Dys1A+/- (n22) and Dys1A-/- (n10) during 30-min exposure to an open field arena. Dys1A-/- show increased locomotion during the first 10 min in the open field (Two-way repeated measure ANOVA, time*genotype interaction: F_10,215_=3.04; p=0.001). **p<0.005 vs Dys1A+/+ at the same time point. **d.** Percentage PPI of the 120dB acoustic startle response displayed by Dys1A+/+ (n10), Dys1A+/- (n15) and Dys1A-/- (n12) littermates. Dys1A-/- have lower pre-pulse intensities compared to Dys1A+/+ mice (Two-way repeated measure ANOVA; genotype: F_2,34_=4.44 p=0.019). *p<0.01 vs Dys1A+/+. **e.** Breakpoint during a food-driven operant behavior test with increasing progressive ratio (PR) displayed by Dys1A+/+ (n12), Dys1A+/- (n13) and Dys1A-/- (n10) littermates. Both Dys1A+/- and Dys1A-/- mice showed lower breakpoints than Dys1A+/+ mice (Two-way repeated measures ANOVA; genotype: F_4,64_=2.8; p=0.032). *p<0.05 vs Dys1A+/+. **f.** Schematic drawing summarizing the behavioral data obtained in Dys1A knockout mice, pointing to a major alteration of basal gangliadependent, but not PFC-dependent, phenotypes.

This further indicates a prominent role for Dys1A in astrocytic functioning.

### Dys1A disruption alters basal ganglia-dependent phenotypes

To identify selective Dys1A-dependent behavioral functions, we next performed in Dys1A knock-out mice a comprehensive battery of tests previously applied to mice with disruption of both Dys1 isoforms ^20, 26, 28, 29^.

In agreement with an initial characterization ^27^, Dys1A+/- and -/- mice were viable with no evident alterations in general health and sensory functions. Similar to this previous study, Dys1A-/- mice presented a slightly hyperactive phenotype compared with Dys1A+/+ littermates (Fig. 2c), as in Dys1 knockout mice ^26, 29^. However, locomotor responses to both acute and sub-chronic amphetamine were not affected by deletion of Dys1A (Supplementary Fig. S2). In contrast to Dys1 knockout mice ^28, 30, 31^, and in agreement with Dys1A mice ^27^, no Dys1A-dependent alterations were evident in different social interaction tests (Supplementary Fig. S2). These results indicate that Dys1A is involved in locomotor activity, but not social interactions.

Dys1 is associated with lower executive function performance in mice, healthy humans, and patients with schizophrenia ^20, 26^. Thus, we tested Dys1A mice in the attentional set-shifting task (ASST), which allows assessment of discrete cognitive executive functions with translational validity to humans ^20, 32^. In contrast to Dys1 knockout mice, Dys1A+/- and -/- mice did not show any deficits in extradimensional set-shifting (EDS) but an alteration in serial reversal learning (Supplementary Fig. S2). EDS alterations imply dopamine dysfunctions in PFC ^32, 33^, while serial reversal learning is linked to dopaminergic tone in striatal regions ^32, 34^. This prompted us to assess behaviors more related to basal ganglia dopamine-related functioning.

Prepulse inhibition (PPI) deficits are consistently linked to overactive dopamine/D2 signaling in basal ganglia ^35–37^. We found a gene-dosage effect for reduced PPI in Dys1A+/- and -/- compared with Dys1A+/+ littermates (Fig. 2d). No Dys1A-dependent effects were evident for acoustic startle responses or body weight (Supplementary Fig. S2), excluding potential confounding factors.

Motivation to receive a reward is another behavioral trait strongly related to dopaminergic functioning within the basal ganglia ^38–40^. We found reduced reward-motivated behavior in Dys1A+/- and -/- compared with Dys1A+/+ littermates (Fig. 2e), when tested in a progressive ratio paradigm designed to assess motivational processes ^41^. No Dys1A-dependent differences were present during acquisition phases (Supplementary Fig. S2), excluding deficits in motor coordination, learning and memory.

Overall, these findings point to a prominent involvement of Dys1A in behavioral phenotypes mediated by dopaminergic signaling in the basal ganglia (Fig. 2f).

### Dys1A disruption alters dopamine/D2 homeostasis in basal ganglia

Dopaminergic signaling in basal ganglia is implicated in locomotion, motivation, serial reversal learning, and PPI ^34, 35, 37^, which are altered in Dys1A knockout mice. Thus, we assessed Dys1A modulation of dopaminergic system in basal ganglia and PFC, as comparison.

We first revealed a relative increased expression of Dys1A in GPe compared to PFC and STR (Fig. 3a). Notably, GPe is a brain region enriched in astrocytes ^10^. In contrast, Dys1C was equally expressed in all regions considered (Fig. 3b).

**Figure 3.**
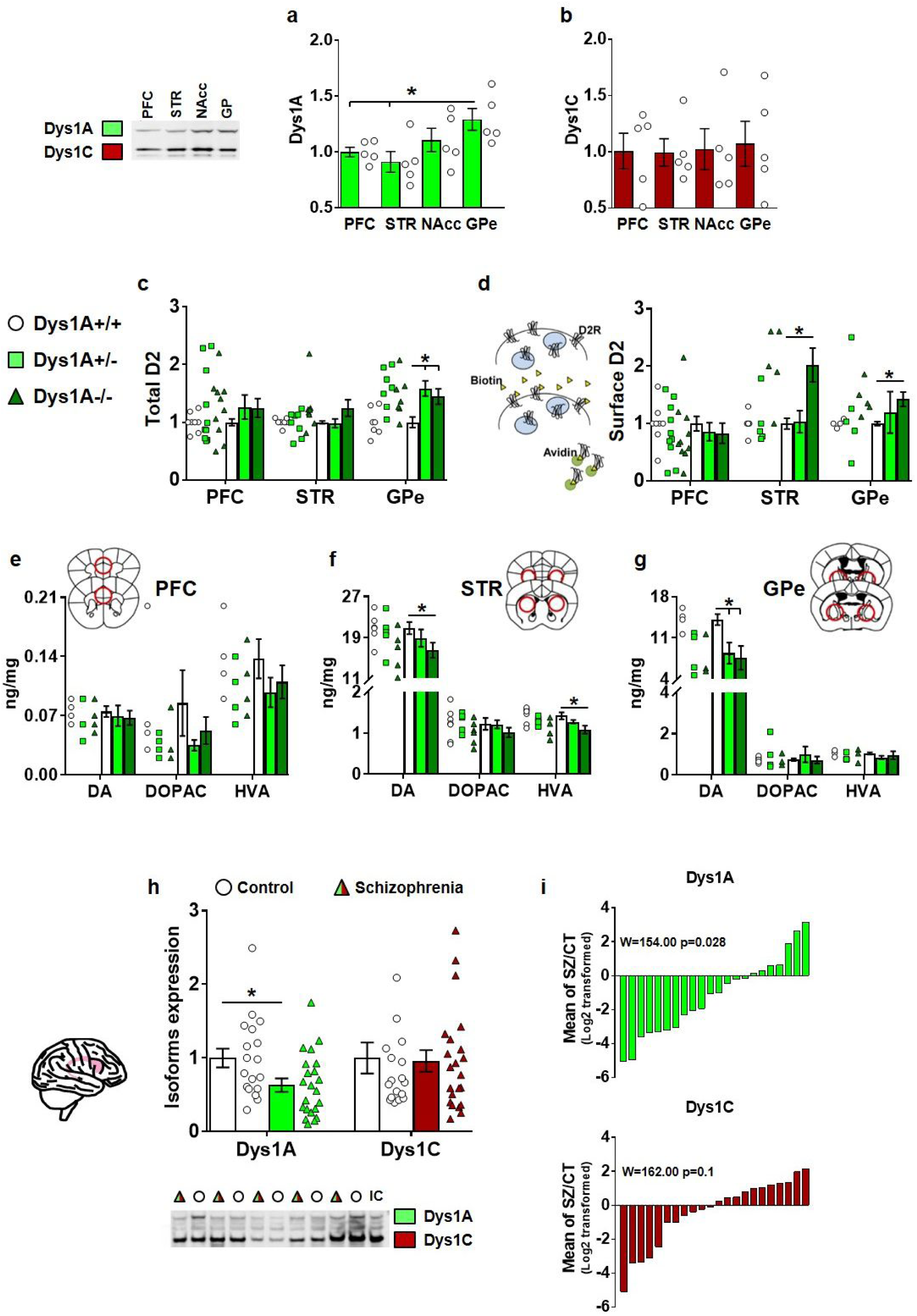
Dys1A alters dopaminergic signaling within the basal ganglia, but not in PFC, and is reduced in the caudate of patients with schizophrenia. Protein expression displayed by C57BL6J adult mice of **a.** Dys1A and **b.** Dys1C isoforms in the PFC, STR, NAcc, and GPe relative to PFC and normalized to their own β-actin. Dys1A expression is higher in GPe compared to PFC and STR (One-way ANOVA: F_3,16_=5.02; p=0.02). *p<0.05 vs PFC and STR. Dys1C expression is uniform across the selected brain areas (One-way ANOVA: F_3,16_=0.045; p=0.98). **c-d.** D2 receptor expression in Dys1A+/+ (n9), Dys1A+/- (n10) and Dys1A-/- (n10). Synaptophysin used as cytoplasmic control, and D2 expression normalized on transferrin as loading control. **c.** Dys1A+/- and Dys1A-/- mice show increased total D2 expression compared to Dys1A+/+ in GPe (One-way ANOVA: F_2,16_=4.93; p=0.021), but not in PFC (F_2,25_=0.724; p=0.49), and STR (One-way ANOVA: F_2,22_=1.44; p=0.26). **d.** Biotinylation protocol, brain slices treated with biotin to label all surface proteins, precipitated by streptavidin. Dys1A-/- mice had increased expression of D2 receptors on cellular surface compared to Dys1A+/+ littermates in STR and GPe (One-way ANOVA: STR F_2,14_=4.20; p=0.04; GPe F_1,6_=6,18; p=0.04), but not in PFC (One-way ANOVA: F_2,24_=0.44; p=0.64). *p<0.05 vs Dys1A+/+. **e-g.** Dopamine (DA), DOPAC, and HVA content by HPLC, expressed as ng/mg of tissue in **e.** PFC, **f.** STR, and **g.** GPe dissected from Dys1A+/+ (n4), Dys1A+/- (n4), and Dys1A-/- (n4) littermates. No Dys1A-dependent changes were observed in PFC (One-way ANOVA: F_2,5_=0.23; p=0.80). Dys1A-/- show reduced DA and HVA levels relative to Dys1A+/+ in the STR (One-way ANOVA: F_1,10_=5.17; p=0.05). Dys1A+/- and Dys1A-/- show reduced DA levels than Dys1A+/+ in GPe (F_1,10_=10.44; p=0.02). *p<0.05 vs Dys1A+/+. **h.** Expression of Dys1A and Dys1C isoforms in postmortem caudate from 22 patients with schizophrenia (Schizophrenia) and 18 matched healthy subjects (Control). No differences were present in non-diagnostic variables (i.e., age, sex, post-mortem interval, pH: Supplementary Fig. S4). Expression of Dys1A, normalized by β-actin, is significantly reduced in the caudate of patients with schizophrenia compared to control subjects (t-test: p=0.02). Dys1C expression is not changed between the two groups (t-test: p=0.71). *p<0.05 vs Control. **i.** Plotting of β-actin normalized data for Dys1A and Dys1C for all case-control pairs. Each bar indicates the log2 transformed ratio of isoform in a schizophrenia case compared to that in its matched control (i.e. the ratio for one case-control pair). Pair-wise analysis of these ratios (Wilcoxon signed-rank test) showed significant difference between schizophrenia cases and their matched controls for Dys1A (W=154.00; p=0.02), but not for Dys1C (W=162.00; p=1.00). Bar graphs show mean ± s.e.m.

Dys1A disruption increased the expression of total D2 receptors in GPe, but not in PFC or STR (Fig. 3c). Moreover, Dys1A disruption increased cellular surface D2 receptors in STR and GPe, but not in PFC (Fig. 3d). Notably, disruption of both Dys1 isoforms resulted in comparable phenotypes in both PFC and striatal regions ^20, 26^ Supplementary Fig. S3). Similarly, a Dys1A genotypic effect on dopamine content was present in STR and GPe, but not in PFC (Fig. 3e-g). In particular, Dys1A-/- mice had lower dopamine levels than Dys1A+/+ in both STR and GPe, and lower HVA levels in STR (Fig. 3f-g). DOPAC/dopamine and HVA/dopamine ratios were indistinguishable across genotypes in all regions, suggesting a normal rate of dopamine catabolism (Supplementary Fig. S3). No Dys1A genotype effects on levels of noradrenaline, serotonin, and 5HIAA were evident (Supplementary Fig. S3).

Thus, consistent with our behavioral assessments, selective disruption of the Dys1A isoform altered the dopaminergic system in basal ganglia, but not in PFC.

### Dys1A is reduced in the caudate of schizophrenia cases

To verify if Dys1A-modulation of basal ganglia-related phenotypes may have clinical relevance, we measured Dys1 isoforms in the caudate of schizophrenia cases and matched healthy controls (Supplementary Fig. S4). This revealed reduced Dys1A, but not Dys1C, in patients with schizophrenia compared with controls (Fig. 3h). Notably, previous findings reported reduced Dys1C, but not Dys1A, in the PFC of patients with schizophrenia ^15^. To directly compare our results to previous reports, we calculated a mean case control ratio where zero indicates no differences between cases and controls, negative values reduced expression in schizophrenia, and positive values increased expression. In the caudate, Dys1A was reduced in 15 out of 22 case-control pairs, while Dys1C ratios were inconsistent in direction, and of generally smaller magnitude (Fig. 3i). Together, these data suggest that Dys1A may have a role in basal ganglia-related schizophrenia pathobiology.

### Selective disruption of Dys1A in astrocytes

To directly assess the specific role of astrocytic Dys1A in basal ganglia dopaminergic and behavioral phenotypes, we next backcrossed Dys1A^flx/flx^ with inducible Glast CreERT2 mice (Fig. 4a). Selective deletion of Dys1A and expression of tdTomato reporter in astrocytes was triggered by tamoxifen injection in adult Dys1AGlast mice, to exclude developmental effects. Employing fluorescence-activated cell sorting (FACS), we isolated tdTomato-positive astrocytes from the striatal region (STR+GPe) of Dys1AGlast mice (Fig. 4b). We confirmed that these cells were enriched in Glast compared to tdTomato-negative cells (Fig. 4c). In agreement with previous reports ^42^, the GlastCre- ERT2 system was very selective for astrocytes, as we found no traces of the NeuN neuronal marker in purified tdTomato-positive cells and equal Glast expression in Dys1AGlast+/+ and -/- mice (Fig. 4d). Importantly, Dys1A expression in Glast-positive astrocytes was abolished in Dys1AGlast-/- mice (Fig. 4e). Compared to Dys1AGlast+/+ littermates, Dys1AGlast-/- mice showed the same GFAP signal (Fig. 4f, and Supplementary Fig. S5), no alteration in astrocytes density (Fig. 4g), and similar astrocytic morphology (Fig. 4h).

**Figure 4.**
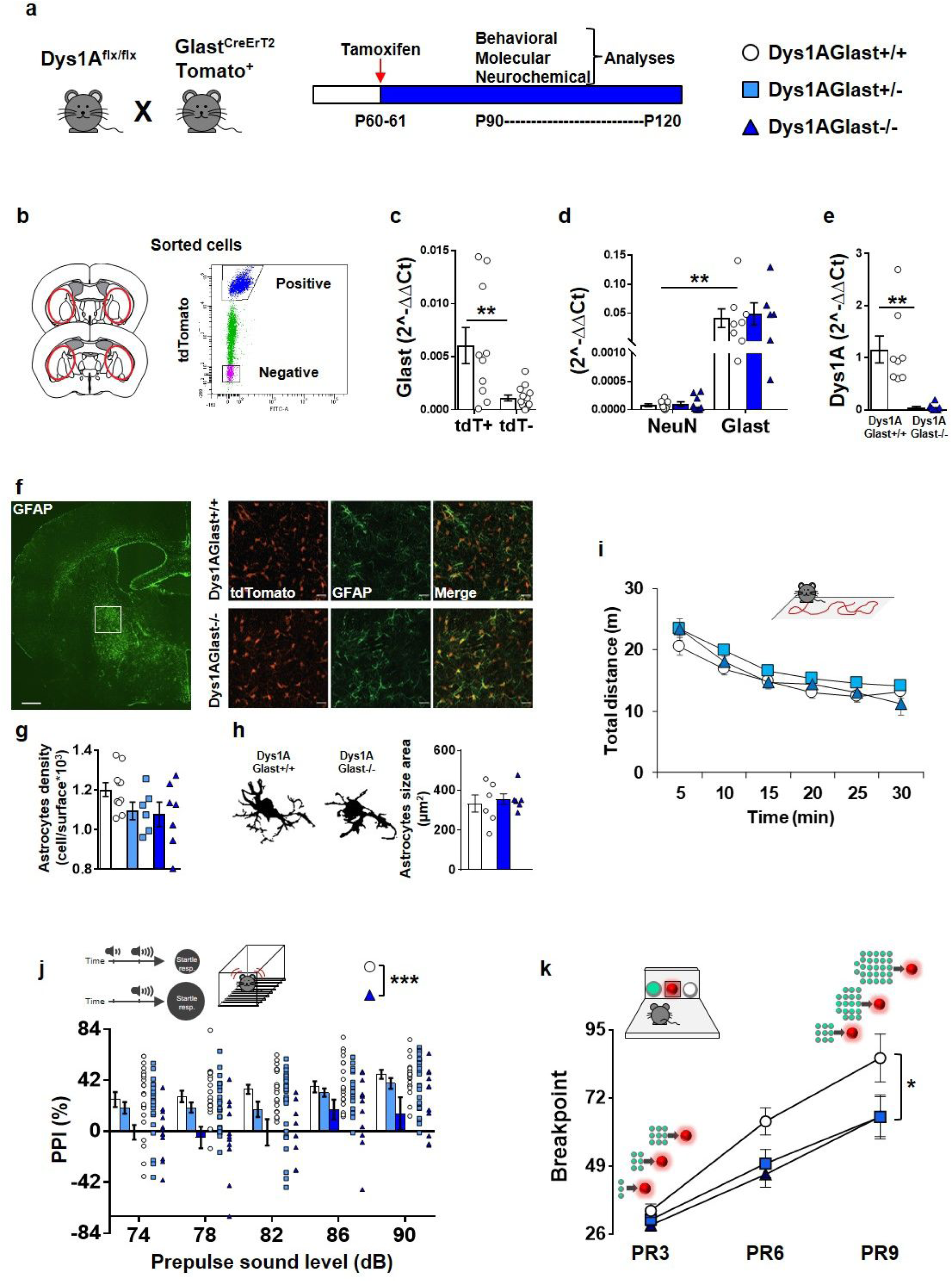
Dys1A disruption in astrocytes is sufficient to induce sensorimotor gating and motivational deficits. **a.** Experimental design and timeline to generate reduction (+/-) or absence (-/-) of Dys1A in astrocytes of adult mice subsequently subjected to molecular and behavioral evaluation. Dys1Afloxed mice were bred with conditional Glast^CreErT2^Tomato^+^ mice, and offspring were treated with tamoxifen at post-natal days 60-61 to then be tested between post-natal days 90-120. **b.** Experimental design, and gating strategy to FACS-sorted astrocytes for subsequent RT-qPCR analyses. **c.** Relative mRNA expression of Glast assessed by RT-qPCR in tdTomato-positive (tdT+) and tdTomato-negative (tdT-) cells sorted from the basal ganglia of Dys1AGlast+/+ (n10) and Dys1AGlast-/- (n13) mice. tdT+ cells show increased expression of Glast compared to tdT- cells (t-test: t=3.25, df=21, *p<*0.005). **p<0.005 tdT+ vs tdT- cells. **d.** Relative mRNA expression of NeuN and Glast assessed by RT-qPCR in tdTomato-positive cells sorted from the basal ganglia of Dys1AGlast+/+ (n10) and Dys1AGlast-/- (n10) mice. tdTomato-positive cells show an equal expression of Glast in Dys1AGlast+/+ and Dys1AGlast-/- mice (Two-way ANOVA; genotype effect: F_1,12_=0.084; p=0.78), but no detectable levels of NeuN (Two-way ANOVA; gene expression: F_1,12_=13.16; p=0.003). **p<0.005 Glast vs NeuN. Expression levels were normalized by Gapdh expression. **e.** Relative mRNA expression of the Dys1A isoform assessed by RT-qPCR in tdTomato-positive cells sorted through FACS from the basal ganglia of Dys1AGlast+/+ (n8) and Dys1AGlast-/- (n8) mice, showing the abolishment of Dys1A expression from Glast-positive astrocytes in the latter group (One-way ANOVA; F_1,14_=18.32; p=0.0008). **p<0.005 vs Dys1AGlast+/+. Expression levels are normalized by Gapdh expression. Data shown as fold-change compared with Dys1AGlast+/+ control mice. **f.** Left: representative 10x confocal images of GFAP-stained brain section at the level of GPe, white square indicates the area magnified 20x for subsequent analyses (scale bar 500µm); right: representative 20x confocal images of GPe brain sections from Dys1AGlast+/+ and -/- mice showing Glast-/tdTomato-positive astrocytes and GFAP-immunoreactivity (scale bar 20µm). **g.** There was no difference in Glast/tdTomato-positive astrocytes density in the GPe (1000 cells x mm^2^) between Dys1AGlast+/+ (10), Dys1AGlast+/- (6), and Dys1AGlast-/- (7) mice (One-way ANOVA: F_2,20_=2.33; p=0.12). **h.** Representative images and quantification of GFAP and Glast-positive astrocytes morphology in the GPe of Dys1AGlast+/+ (n6) and Dys1AGlast-/- (n6) mice. No genotype-dependent difference was observed in the astrocytic surface area measured by GFAP immunoreactivity (t-test: t_10_=-0.46, p=0.65). **i.** Spontaneous distance traveled by Dys1AGlast+/+ (n20), Dys1AGlast+/- (n24), and Dys1AGlast-/- (n11) during 30 minutes exposure to an open field arena. No genotype differences were evident (Two-way repeated measure ANOVA, genotype effect: F_2,52_=2.79; p=0.07; time*genotype effect: F_10,260_=0.91; p=0.53). **j.** Percent pre-pulse inhibition (PPI) of the 120dB acoustic startle response displayed by Dys1AGlast+/+ (n21), Dys1AGlast+/- (n24), and Dys1AGlast-/- (n13) littermates. Dys1AGlast-/- have reduced PPI compared to Dys1AGlast+/+ mice (Two-way repeated measure ANOVA; genotype effect: F_2,54_=8.59 p=0.0006). ***p<0.0005 vs Dys1AGlast+/+. **k.** Breakpoint during a food-driven operant behavior test with increasing progressive ratio (PR) displayed by Dys1AGlast+/+ (n15), Dys1AGlast+/- (n15), and Dys1AGlast-/- (n9) littermates. Both Dys1AGlast+/- and Dys1AGlast-/- mice showed lower breakpoints than Dys1AGlast+/+ mice (Two-way repeated measure ANOVA; genotype effect: F_2,36_=4.35; p=0.020). *p<0.05 vs Dys1AGlast+/+. Bar and line graphs show mean ± s.e.m.

These data demonstrate a selective inducible deletion of the Dys1A in astrocytes, with no major alterations in astrocytic anatomy.

### Dys1A disruption in astrocytes induces motivational and sensorimotor gating deficits

To investigate the role played by astrocytic Dys1A in behavioral functions, we tested Dys1AGlast+/+, +/- and -/- mice for phenotypes that were altered by Dys1A disruption in all cell types (Fig. 2).

No Dys1AGlast genotype effect was evident in exploratory activity (Fig. 4i). Similar to ubiquitous Dys1A disruption, selective Dys1A deletion in astrocytes was sufficient to impair PPI sensorimotor gating abilities (Fig. 4j), while having no effects on startle reactivity or body weight (Supplementary Fig. S5). Furthermore, as observed in Dys1A knockout mice (Fig. 2), Dys1AGlast+/- and -/- mice showed reduced motivation to work for a rewarding stimulus (Fig. 4k), while no Dys1AGlast genotype effects were evident on the acquisition of this operant task (Supplementary Fig. S5).

These results indicate a specific role for astrocytic Dys1A in modulating basal ganglia-related sensorimotor gating and motivational processes.

### Dys1A disruption in astrocytes alters basal ganglia dopamine homeostasis and increases D2 in astrocytes

We then investigated the effects of astrocytic Dys1A on dopaminergic signaling in basal ganglia.

Similar to ubiquitous Dys1A disruption (Fig. 3), astrocytic Dys1A disruption reduced dopamine levels primarily in GPe and marginally in STR (Fig. 5a). No effects were observed in PFC (Supplementary Fig. S5). No Dys1AGlast effect was evident for DOPAC levels (Fig 5b), while HVA levels were reduced in Dys1AGlast knockout mice compared with wild-type littermates in both STR and GPe (Fig. 5c). These findings demonstrate that disruption of Dys1A in astrocytes is sufficient to reduce dopamine levels, primarily in GPe.

**Figure 5.**
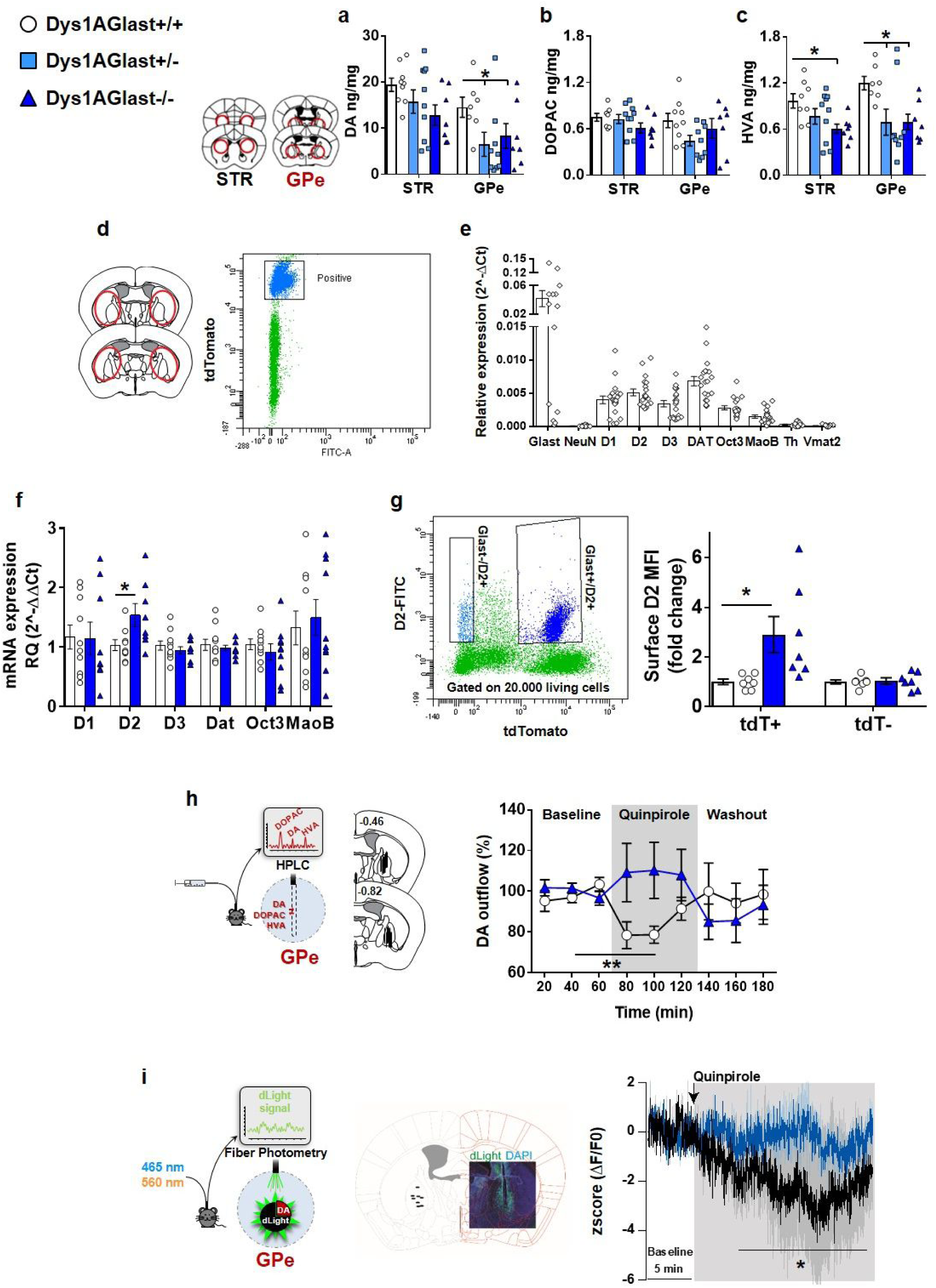
Dys1A disruption in astrocytes alters dopamine/D2 homeostasis in basal ganglia. **a.** Dopamine (DA), **b.** DOPAC, and **c.** HVA content by HPLC analyses, expressed as ng/mg of dissected STR or GPe tissues displayed by Dys1AGlast+/+ (n9), Dys1AGlast+/- (n10), and Dys1AGlast-/- (n7) littermates. In the STR, Dys1AGlast-/- had reduced HVA levels (One-way ANOVA: F_2,22_=3.52; p=0.04), but no alterations in DA (One-way ANOVA: F_2,23_=2.15; p=0.14), and DOPAC levels (One-way ANOVA: F_2,21_=1.40; p=0.27), compared to Dys1AGlast+/+ mice. In the GPe, Dys1AGlast-/- had reduced DA (One-way ANOVA: F_2,21_=5.34; p=0.03), and HVA levels (One-way ANOVA: F_2,20_=4.46; p=0.02), but no alterations in DOPAC (One-way ANOVA: F_2,22_=2.08; p=0.15), compared to Dys1AGlast+/+ mice. *p<0.05 vs Dys1AGlast+/+ mice. **d.** Gating strategy to FACS-sorted astrocytes from STR+GPe regions for subsequent RT-qPCR analyses for astrocytic dopaminergic markers. **e.** Relative mRNA expression of Glast, NeuN, dopamine receptors (D1, D2, D3), Dat, Oct3, MaoB, TH, and Vmat2 assessed by RT-qPCR in tdTomato-positive cells sorted from basal ganglia (STR+GPe) of Dys1AGlast+/+ and Dys1AGlast-/- mice (n20). tdTomato-positive cells show no detectable levels of NeuN, Th and Vmat2. Expression levels are normalized by Gapdh mRNA expression. **f.** Relative mRNA expression of dopamine receptors (D1, D2, D3), Dat, Oct3 and MaoB, assessed by RT-qPCR in tdTomato-positive cells sorted from basal ganglia of Dys1AGlast+/+ (n10) and Dys1AGlast-/- (n10) littermates, revealing an increased astrocytic D2 receptor expression in Dys-Glast-/- mice (t-tests: t_17_=-2.42, p*=*0.029), and no genotype effect for the other assessed makers (D1: t_17_=0.07, p*=*0.95; D3: t_17_=0.81, p*=*0.43; Dat: t_17_=0.50, p*=*0.62; Oct3: t_17_=0.76, p*=*0.46; Maob: t_17_=-0.00, p=1.00). Data shown as fold-change compared with Dys1AGlast+/+ control mice. *p<0.05 vs Dys1AGlast+/+. **g.** Flow cytometry gating strategy and surface D2 receptor protein expression quantification in tdTomato-positive (tdT+) and negative (tdT-) cells from the basal ganglia of Dys1AGlast+/+ (n7) and Dys1AGlast-/- mice (n7) expressed as mean fluorescence intensity (MFI), revealing increased astrocytic surface D2 expression in DysGlast-/- mice compared to Dys1AGlast+/+ only in tdTomato-positive cells (Two-way ANOVA: genotype*cell-type interaction: F_1,12_=5.71; p=0.03). Data shown as fold-change compared with Dys1AGlast+/+ control mice. *p<0.01 Dys1AGlast-/- tdT+ vs all other groups. **h.** Schematic of the experiment and localization of probe dialyzing portion within the GPe. Dys1AGlast+/+ (n11) and Dys1Glast-/- (n9) littermates were implanted with a dialysis probe for measurement of basal extracellular dopamine levels and quinpirole-induced dopamine release. Quinpirole infusion (gray area) decreased extracellular dopamine release in the GPe of Dys1AGlast+/+ mice, but not in Dys1AGlast-/- littermates (Two-way RM ANOVA: genotype*time interaction, F_8,148_=2,18; p<0.05). **p<0.005 Dys1AGlast+/+ quinpirole vs baseline. **i.** Schematic of the fiber photometry experiment and representative post-hoc localization of the tip of the optic fiber within GPe (left) and low magnification confocal images of the viral spread and optic fiber placement (right). Dys1AGlast+/+ (n4) and Dys1Glast-/- (n4) littermates were first injected with the adeno-associated virus (AAV) to express dLight and successively implanted with an optic fiber for recording of synaptic release of dopamine using fiber photometry. Quinpirole (0.5 mg/kg, i.p.) decreased dLight signal compared to the 5-min baseline period before injection in the GPe of Dys1AGlast+/+ mice, but not in Dys1AGlast-/- littermates (Two-way RM ANOVA: geno-type*time interaction, F_8,148_=34.32; p<0.0005). *p<0.05 Dys1AGlast+/+ quinpirole vs their baseline.

We next assessed the impact of Dys1A disruption in astrocytes on the expression of genes involved in dopaminergic signaling and metabolism in purified astrocytes isolated from the basal ganglia region (Fig. 5d). We found detectable levels of mRNA expression for dopamine receptors D1, D2, and D3, for the plasma membrane dopamine (DAT) and organic cation 3 (OCT3) transporters, and the metabolic enzyme monoamine oxidase type B (MAOB (Fig. 5e). No detectable signals were evident for tyrosine hydroxylase (TH), the rate-limiting enzyme in dopamine synthesis, and for the vesicular monoamine transporter VMAT2 that has been implicated in astrocytic dopamine homeostasis in PFC^3^. We excluded neuronal contamination, as no detectable levels of the NeuN marker were found (Fig. 5e).

Among all these markers, only astrocytic expression of dopamine D2 receptor was altered, with Dys1AGlast-/- mice showing higher D2 mRNA level than Dys1AGlast+/+ (Fig. 5f). This is in accordance with and extends the effects of ubiquitous disruption of Dys1 on expression of D2 receptors in basal ganglia (Supplementary Fig. S6). Similarly, in drosophila, ubiquitous, glia- or neuron-specific disruption of dDys equivalently increased D2 expression (Supplementary Fig. S6). Notably, we confirmed by flow cytometry an increased protein expression of D2 receptors on the surface of Glast/tdTomato positive astrocytes from Dys1AGlast-/- mice compared with Dys1AGlast+/+ (Fig. 5g). In these same mice, no genotype-dependent differences in D2 protein expression were evident on Glast/tdTomato negative cells (Fig. 5g), confirming the selective upregulation of D2 receptor in astrocytes upon Dys1A disruption, without any compensatory D2 perturbation in other cells.

These data suggest that Dys1A-dependent behavioral and dopaminergic alterations might relate to increased astrocytic D2 mechanisms.

### Dys1A disruption in astrocytes alters D2-dependent dopamine dynamics in GPe

We then investigated the impact of astrocytic Dys1A deficiency on D2-mediated dopamine modulation in GPe *in vivo*.

Local GPe perfusion by reverse microdialysis of the dopamine D2-like receptor agonist quinpirole reduced dopamine outflow in Dys1AGlast+/+ mice (Fig. 5h, Supplementary Fig. S5). Similarly, systemic injection of quinpirole reduced dopamine binding in Dys1AGlast+/+ mice (Fig. 5i), as revealed by the dopamine-sensor dLight fiberphotometry ^43^. In contrast, both indices of D2-mediated dopamine modulation were lost in Dys1AGlast-/- mice (Fig. 5h, i). The effect of amphetamine on dopamine binding was unaltered by astrocytic Dys1A deletion (Supplementary Fig. S5), in line with behavioral findings (Supplementary Fig. S2), and strengthening a preferential link of astrocytic Dys1A with D2 functioning.

Overall, these data demonstrate that astrocytic D2 overexpression, upon astrocytic Dys1A disruption, is associated with loss of D2-mediated inhibition of dopamine release in basal ganglia. This extends other evidence implicating astrocytes in the inhibitory control of dopamine release within the basal ganglia ^10, 44^.

### Dys1A disruption in astrocytes delays reward-expectancy dopamine dynamics in GPe

To directly correlate behavioral functioning with dopamine dynamics within the GPe to, and its modulation by astrocytic Dys1A, we used the dLight dopamine sensor during progressive ratio testing (Fig. 6a).

**Figure 6.**
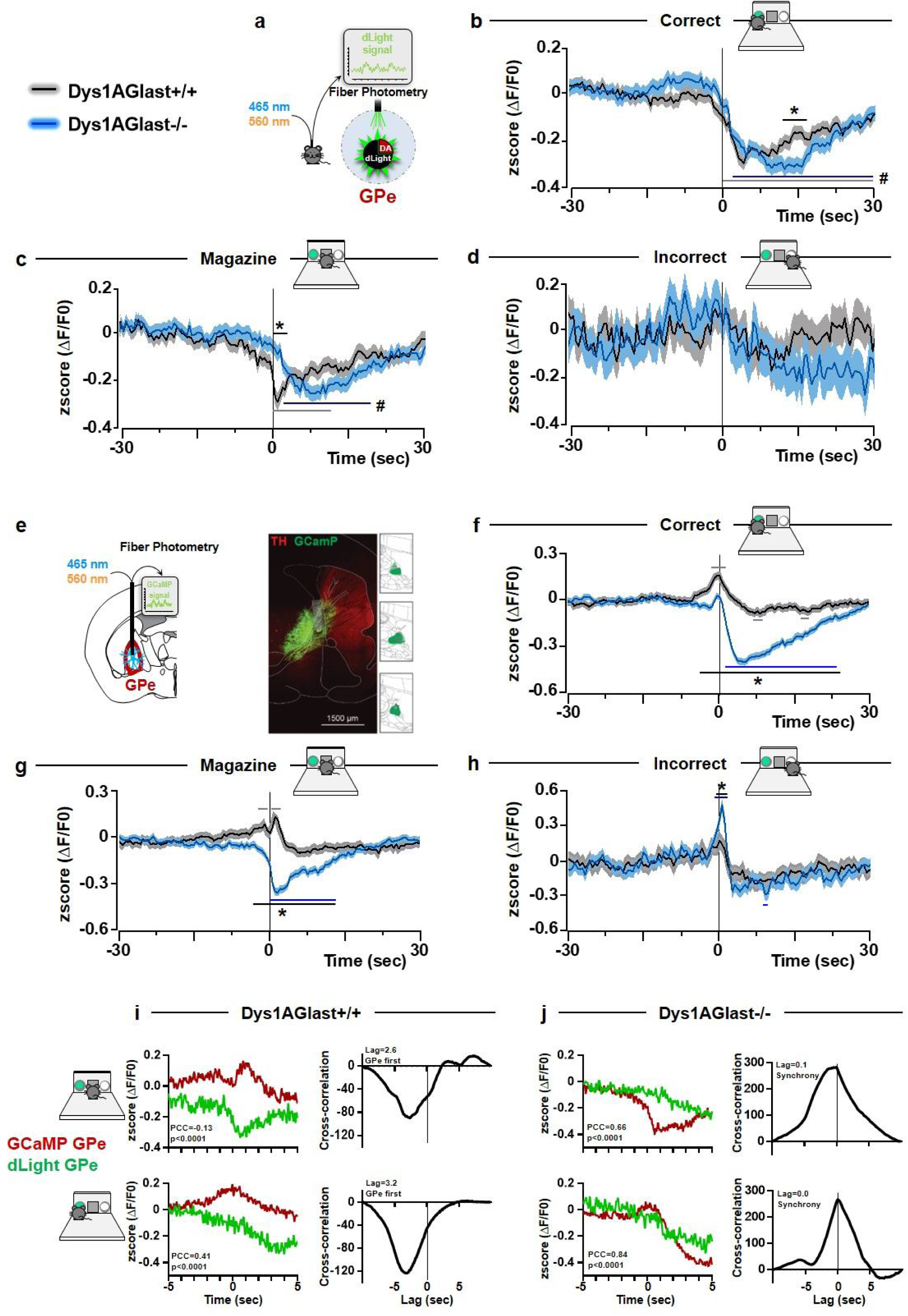
Dys1A disruption in astrocytes alters astrocytes-dopamine communication related to reward-expectancy in GPe. **a.** Schematic of the fiber photometry experiment performed in Dys1AGlast+/+ (n4) and Dys1Glast-/- (n4) littermates injected with the AAV to express dLight and successively implanted with an optic fiber for recording of synaptic release of dopamine in the GPe during the progressive ratio test. **b**. Analyses of the dLight signal during responses to correct nose pokes (Correct) revealed a ΔF/F0 decrease compared to baseline values in the GPe of both Dys1AGlast+/+ (from 0 to 29 sec after a correct poke) and Dys1AGlast-/- mice (from 2 to 30 sec after a correct poke), and decreased values in Dys1AGlast-/- compared to Dys1AGlast+/+ from 13 to 17 seconds after a correct poke (Two-way RM ANOVA, genotype-by-time interaction: F_120,253080_=4.41; p<0.0001). #p<0.05 vs own genotype baseline. *p<0.05 vs Dys1AGlast+/+ at the same time interval. **c.** Analyses of the dLight signal in the GPe during food magazine entrance revealed a decreased (ΔF/F0) signal in Dys1AGlast+/+ from the entrance to the magazine up to 10 sec later, and from 3 sec up to 21 in Dys1AGlast-/- mice. #p<0.05 vs own genotype baseline. Moreover, compared to Dys1AGlast+/+ littermates, Dys1AGlast-/- mice had higher dLight signal from 0 to 2 sec following magazine entrance *p<0.05 vs Dys1AGlast+/+ at the same time interval. (Two-way RM ANOVA, genotype-by-time interaction: F_120,249240_=4.61; p<0.0001). **d.** Analyses of the dLight signal in the GPe during incorrect nose pokes (Incorrect) revealed no ΔF/F0 changes compared to baseline values of both Dys1AGlast+/+ and Dys1AGlast-/- mice nor genotype-dependent effects (Two-way RM ANOVA, genotype-by-time interaction: F_120,35400_=1.96; p<0.0001). **e.** Experimental design to monitor astrocytes Ca^2+^ levels in GPe using a fiber-photometry system in freely behaving Dys1AGlast+/+ (n6) and Dys1AGlast-/- (n5) during the progressive ratio test. Reconstruction of viral expression and location of viral injection sites. Placements and representative images of viral expression (GCamP) and staining for dopaminergic neurons (TH) in the GPe and SNc/VTA in the same mouse. **f.** Astrocytes responses to correct nose pokes (Correct) revealed in Dys1AGlast+/+ mice an increased ΔF/F0 signal from 1 sec before up to 2 sec following the entrance into the correct poke, and decreased signal from 6.5 to 10 sec and from 17 to 19 sec after the correct poke. In Dys1AGlast-/- mice, decreased ΔF/F0 Ca^2+^ signal was evident from 2 up to 25 sec following entrance into the correct poke. (Two-way RM ANOVA, genotype-by-time interaction: F_120,671880_=42.89; p<0.0001). *p<0.05 gray line: Dys1AGlast+/+ vs their own baseline; blue line: Dys1AGlast-/- vs their own baseline; black line: Dys1AGlast-/- vs Dys1AGlast+/+ at the same time interval. **g.** Astrocytes responses to magazine entrance (Magazine) revealed in Dys1AGlast+/+ mice an increased ΔF/F0 signal from 2.5 to 1 sec before and from 1 to 3 sec after magazine entrance. In Dys1AGlast-/- mice, decreased ΔF/F0 Ca^2+^ signal was evident from 0 up to 14 sec following entrance into the magazine (Two-way RM ANOVA, genotype-by-time interaction: F_120,460440_=21.44; p<0.0001). *p<0.05 gray line: Dys1AGlast+/+ vs their own baseline; blue line: Dys1AGlast-/- vs their own baseline; black line: Dys1AGlast-/- vs Dys1AGlast+/+ at the same time interval. **h.** Astrocytes responses to incorrect nose pokes (Incorrect) revealed no effects in Dys1AGlast+/+ mice. In Dys1AGlast-/- mice, increased ΔF/F0 Ca^2+^ signal was evident from -1 to 1 sec following entrance into the incorrect choice, and a decrease from 9 to 10 sec after (Two-way RM ANOVA, genotype-by-time interaction: F_120,73560_=1.75; p<0.0001). *p<0.05 gray line: Dys1AGlast+/+ vs their own baseline; blue line: Dys1AGlast-/- vs their own baseline; black line: Dys1AGlast-/- vs Dys1AGlast+/+ at the same time interval. **i.** Correlation of overall activity (left) and cross-correlation (right) in Dys1AGlast+/+ mice between the GCaMP signal in GPe and dLight in GPe, referred to 5 sec before and 5 sec after magazine (upper) and correct entry (lower). **j.** Correlation of overall activity (left) and cross-correlation (right) in Dys1AGlast-/- mice between the GCaMP signal in GPe and dLight in GPe, referred to 5 sec before and 5 sec after magazine (upper) and correct entry (lower).

Decreased dopamine signal was evident in Dys1AGlast+/+ immediately after a correct poke, while this decrease was delayed of 2 seconds in Dys1AGlast-/- mice (Fig. 6b). Similarly, a sharp decrease of dopamine signal was evident in Dys1AGlast+/+ immediately after entry into the food magazine (Fig. 6c), regardless of presence of a food pellet (Supplementary Fig. S7). In contrast, Dys1AGlast-/- mice showed a more gradual decrease within 3 seconds after magazine entry (Fig. 6c; Supplementary Fig. S7). No behavioral- or genotype-dependent effects on GPe dopamine dynamics were evident for incorrect responses (Fig. 6d). No behavioral-dependent variations on tdTomato control signal were detected (Supplementary Fig. S7).

These results indicate that reward expectancy is associated with decreased dopamine signaling in GPe, and that disruption of Dys1A in astrocytes, which impairs motivation, delays this effect.

### Dys1A disruption in astrocytes alters reward-related astrocytic responses in GPe

To visualize the involvement of GPe astrocytes in motivational behaviors, and the impact of astrocytic Dys1A disruption, we monitored astrocyte population Ca^2+^ levels in GPe in Dys1AGlast+/+ and Dys1AGlast-/- mice while performing the progressive ratio test (Fig. 6e).

Analyses of behaviorally linked Ca^2+^ signaling revealed that astrocytic responses in GPe of Dys1AGlast+/+ mice increased during a correct choice and food magazine entry and decreased starting 3 seconds after these actions (Fig. 6f,g). No significant Ca^2+^ modulation was evident for incorrect responses in GPe of Dys1AGlast+/+ mice (Fig. 6h). In contrast, Dys1AGlast-/- mice showed a strong decrease of astrocytes Ca^2+^ responses in GPe following correct choices and magazine entrance compared with Dys1AGlast+/+ mice (Fig. 6f,g). Conversely, an increased astrocyte Ca^2+^ response in GPe was evident when an incorrect nose poke was made (Fig. 6h). No effects were evident for GPe astrocytic Ca^2+^ signal when traces were phase-randomized, and for the tdTomato control signal (Supplementary Fig. S7).

This demonstrates an aberrant reduction of reward-related astrocytic responses in GPe upon deletion of Dys1A in astrocytes. This effect could be related to increased astrocytic D2 in Dys1AGlast-/- mice, as D2 stimulation was reported to decrease astrocytes Ca^2+^ responses in GPe and midbrain ^10, 45^.

### Dys1A disruption in astrocytes switches reward-related dopamine-astrocytic communication

In an attempt to relate *in vivo* dopamine with astrocytic dynamics, we quantified reward-related correlations (Pearson’s correlation coefficient, PCC), and timescale cross-correlation between GPe astrocytic activity and GPe dopamine dynamics.

In Dys1AGlast+/+, astrocytes responses in GPe were anti-correlated to GPe dopamine with a lag of 2-3 sec, suggesting that astrocytes responses preceded dopamine fluctuations (Fig. 6i). This is in line with the idea of astrocytes contributing to the inhibition of dopamine release ^10, 44^. Conversely, Dys1AGlast-/- show a strikingly high positive correlation paired with a precise synchrony (lag=0.1- 0.0sec) between astrocyte activity and dopamine dynamic (Fig. 6j). This strengthens the evidence of an impaired astrocytic inhibitory control over dopamine release following Dys1A disruption in astrocytes, which could then result in a delayed reward-related dopamine decrease.

These data support the conclusion that Dys1A disruption in astrocytes reverts reward-related astrocyte-to-dopamine communication in GPe.

### D2 disruption in GPe astrocytes increases motivation and GPe dopamine levels

To test if D2 receptors in astrocytes are responsible for astrocytic Dys1A modulation of basal ganglia behavioral and dopaminergic processes, we selectively deleted D2 receptors in GPe astrocytes (GFAP-D2-GP-/-; Fig. 7a).

**Figure 7.**
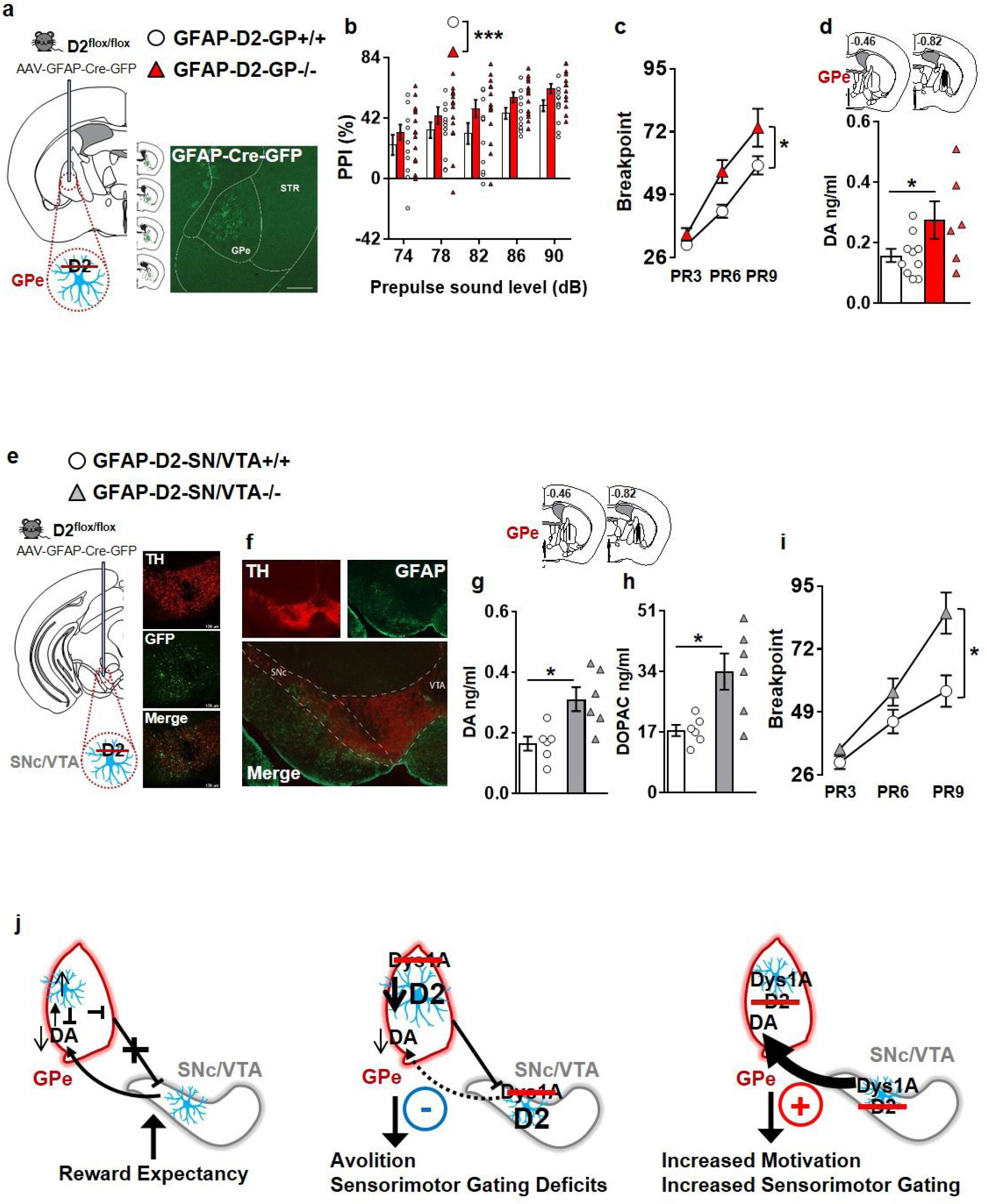
D2 disruption in astrocytes in GPe or SNc/VTA increased PPI, motivation and dopamine outflow within the GPe. **a.** Experimental design to selectively delete dopamine D2 receptors of D2^flox/flox^ mice restricted to GPe-astrocyte, bilaterally. Reconstruction of viral spread across the GPe along anteroposterior axis. Low magnification images of a representative viral spread on a coronal slices of a D2^flox/flox^ mice injected with AAV-GFAP-Cre-GFP (scale bar=500µm). Findings were replicated in two independent experiments with similar results. **b.** Percentage PPI of the 120dB acoustic startle response displayed by control mice bilaterally injected in the GP with the AAV-GFAP-Cre-GFP (GFAP-D2-GP+/+, n12), and D2^flox/flox^ littermates bilaterally injected in the GPe with the AAV-GFAP-Cre-GFP (GFAP-D2-GP-/-, n14). GFAP-D2-GP-/- have increased PPI compared to GFAP- D2-GP+/+ mice (Two-way repeated measure ANOVA; genotype effect: F_1,25_=4.70 p=0.040). *p<0.05 vs GFAP-D2-GP+/+. **c.** Breakpoint during a food-driven operant behavior test with increasing progressive ratio (PR) displayed by GFAP-D2-GP+/+ (n12), and GFAP-D2-GP-/- (n12) littermates. GFAP-D2-GP-/- mice showed higher breakpoints than GFAP-D2-GP+/+ mice (Two-way repeated measure ANOVA; genotype effect: F_1,22_=5.78; p=0.025). *p<0.05 vs GFAP-D2-GP+/+. **d.** GFAP-D2-GP+/+ (n11) and GFAP-D2-GP-/- (n8) littermates were implanted with a dialysis probe for measurement of basal extracellular dopamine levels. Top panel show the localization of probe dialyzing portion within the GPe. Three mice were excluded for misplaced probe. GFAP-D2-GP-/- show higher extracellular dopamine levels compared to GFAP-D2-GP+/+ mice (t-test: t_14_=-2.13, p*=*0.05). *p<0.05 vs GFAP-D2-GP+/+. **e.** Experimental design to selectively delete dopamine D2 receptors in the astrocytes of adult D2^flox/flox^ mice only within SNc/VTA, bilaterally. Example images of SNc/VTA stained for tyrosine hydroxylase (TH, to visualize the dopaminergic neurons) and for GFP (to identify the cells infected by the virus). **f.** Distribution of GFAP-positive astrocyte in the VTA/SN section co-stained for TH as a maker for the dopaminergic neurons. **g-h.** GFAP-D2- VTA/SN+/+ (n6) and GFAP-D2-SN/VTA-/- (n6) littermates were implanted with a dialysis probe for measurement of basal extracellular dopamine levels in the GPe (top figure shows the localization of probes dialyzing portion within the GPe). GFAP-D2-SN/VTA-/- showed increased basal extracellular **g.** dopamine (t-test: t_10_=-3.20, p*=*0.009), and **h.** DOPAC levels (t-test: t_10_=-3.10, p*=*0.011) than GFAP-D2-SN/VTA+/+ littermates. *p<0.05 vs GFAP-D2-SN/VTA+/+. **i.** Breakpoint during a food-driven operant behavior test with increasing progressive ratio (PR) displayed by GFAP-D2- SN/VTA+/+ (n10), and GFAP-D2-SN/VTA-/- (n14) littermates. GFAP-D2-SN/VTA-/- mice showed higher breakpoints than GFAP-D2-SN/VTA+/+ mice (Two-way repeated measure ANOVA; geno-type effect: F_1,22_=5.28; p=0.031). *p<0.05 vs GFAP-D2-SN/VTA+/+. **j.** Schematic figure model depicting astrocytes-dependent Dys1A/D2 signaling pathways involved in basal ganglia dopamine-related modulation of motivational and sensorimotor gating abilities.

Opposite to astrocytic Dys1A disruption (Fig. 4), D2 disruption in GPe astrocytes increased sensorimotor gating abilities (Fig. 7b), while not affecting startle reactivity or body weight (Supplementary Fig. S8). Opposite to astrocytic Dys1A disruption (Fig. 4), GFAP-D2-GP-/- mice showed increased motivation to work for a reward compared to GFAP-D2-GP+/+ (Fig. 7c), while no GFAP- D2-GP genotype effects were evident in the acquisition of the basic operant task (Supplementary Fig. S8). No differences were evident between GFAP-D2-GP-/- and GFAP-D2-GP+/+ in exploratory activity (Supplementary Fig. S8). Finally, opposite to astrocytic Dys1A disruption (Fig. 5), GFAP-D2- GP-/- mice showed increased dopamine levels in GPe (Fig. 7d). No differences between GFAP-D2- GP-/- and GFAP-D2-GP+/+ mice were evident in levels of DOPAC or HVA (Supplementary Fig. S8).

Overall, these results demonstrate a direct link between GPe astrocytic D2 levels, GPe dopamine levels, sensorimotor gating and motivational processes. In particular, increased D2 in astrocytes in the basal ganglia is associated with impaired PPI, reduced motivation and reduced dopamine levels in GPe. Conversely, decreased astrocytic D2 in GPe increased PPI, motivation, and GPe dopamine levels.

### D2 disruption in SNc/VTA astrocytes increases motivation and GPe dopamine levels

Dopamine in GPe is released by midbrain dopaminergic neurons ^46, 47^, whose activity can be modulated by astrocytes ^48^. As in STR/GPe (Fig. 5), Dys1A disruption in astrocytes increased D2 astrocytic expression also in the midbrain (Supplementary Fig. S8). Thus, we finally tested if midbrain astrocytic D2 signaling can also influence basal ganglia-dependent behavioral and dopaminergic pheno-types.

We deleted D2 receptors in astrocytes within the substantia nigra pars compacta (SNc) and ventral tegmental area (VTA) regions (GFAP-D2-SN/VTA-/-; Fig. 7e). GFAP expression was more pronounced in SNc compared to VTA (Fig. 7f). This might result in a bias towards greater D2 deletion in astrocytes within SNc, the major source of dopaminergic projections to GPe ^46, 47^. Notably, opposite to astrocytic Dys1A disruption, and similar to astrocytic D2 disruption in GPe, GFAP-D2-SN/VTA-/- mice showed increased dopamine and DOPAC levels in GPe (Fig. 7g,h). No difference between these two groups was evident in HVA levels (Supplementary Fig. S8). Opposite to astrocytic Dys1A disruption, and similar to astrocytic D2 disruption in GPe, GFAP-D2-SN/VTA-/- mice showed increased motivation to work for a reward compared to GFAP-D2-SN/VTA+/+ (Fig. 7i). No genotype effects were evident in acquisition of the operant task (Supplementary Fig. S8). No genotype differences were evident in exploratory activity and body weight (Supplementary Fig. S8). These results demonstrate a direct link between SNc/VTA astrocytic D2, dopamine levels within the GPe, and motivational behavior.

Overall, these data clarify that astrocytic D2 receptors can cause behavioral and basal ganglia dopamine alterations by a local effect on GPe and through the modulation of dopaminergic transmission from the SNc/VTA.

## Discussion

In this study, we reveal Dys1A as a molecule participating in astrocytic modulation of behavioral and dopaminergic/D2 pathways within the basal ganglia, with implications for schizophrenia. Notably, the Dys1A isoform which is reduced in the caudate of patients with schizophrenia, is the only Dys1 isoform expressed by astrocytes. Disruption of Dys1A in astrocytes is sufficient to alter dopamine/D2 dynamics within basal ganglia and, accordingly, induce basal ganglia-dependent reward and sensorimotor gating deficits. Finally, we demonstrate that these behavioral and dopaminergic alterations depend on astrocytic D2 receptors levels within GPe and SNc/VTA regions (Fig. 7j).

Dys1A affected basal ganglia dopaminergic and behavioral functions, while sparing these processes within the cortex. The differential control by astrocytic Dys1A of dopamine-related behaviors between cortical and basal ganglia regions adds to increasing evidence for heterogeneous astrocytemediated processes across different brain regions ^12–14^. In contrast to Dys1A, the expression of Dys1C increases during cortical development, and is selectively reduced in PFC of patients with schizophrenia ^17^. This suggests that Dys1-dependent dopaminergic and behavioral alterations at the cortical level ^20, 22, 26, 49^ might be more related to neuronal Dys1C. Most importantly, our findings uncovers a novel astrocytic mechanism as a player in the dopaminergic cortical/basal ganglia dichotomy and related behavioral abnormalities consistently reported in schizophrenia ^50, 51^.

We reveal that the interaction between astrocytes and dopamine system at the level of basal ganglia has major effects on motivated behavior and saliency detection. In particular, Dys1A reduction in astrocytes increased levels of astrocytic D2 receptors in basal ganglia, and resulted in avolition and sensorimotor gating deficits. Similarly, striatal overexpression of D2 receptors by the CamKII promoter ^52^, which is also present in astrocytes ^53^, reduced motivation ^38^. Conversely, D2 reduction in astrocytes in either GPe or SNc/VTA, resulted in increased motivation and sensorimotor gating abilities. In contrast, both astrocytic manipulations of Dys1A and D2 spared gross motor functions, despite altering dopamine signaling within GPe. Historically, the GPe has been implicated in movement control ^54^. However, in agreement with our findings, recent studies are showing GPe involvement in motivational and sensorimotor gating abilities ^55–58^. Moreover, in agreement with our findings, dopaminergic manipulations within GPe may not alter motor functions ^55, 58, 59^, and D2 receptors do not contribute to amphetamine-evoked astrocytic and locomotor responses ^5^. In contrast, astrocyte-dopamine interactions within VTA-NAcc dopamine/D1 pathways are involved in locomotor processes ^5^. Thus, we here extend previous findings by showing that astrocytes regulate motivational and sensorimotor gating processes by dopamine/D2-dependent mechanisms involving SNc/VTA to STR/GPe pathways. Overall, astrocytic Dys1A and D2 receptors could be cell-specific targets for the treatment of motivational and other neuropsychiatric disorders associated with disrupted dopamine/D2 signaling.

In agreement with increasing evidence ^3, 5, 9, 10, 44^, our findings strengthen the direct implication of astrocytes as modulators of dopaminergic signaling. In particular, we demonstrate a previously unknown contribution of Dys1A in the astrocytic machinery controlling dopamine/D2 levels and functions, especially within GPe. Dopamine in GPe is mainly released through volume transmission ^47^, which may better relate to astrocytic activity ^44, 60^. Conditions of reduced dopamine in GPe and STR are associated with increased astrocytic activity and augmented astrocyte-related GABA inhibition of dopamine release ^10, 44^. In agreement, in physiological conditions (wild-type mice, Fig. 7j), reward expectancy-related decrease of dopamine levels in GPe preceded and was anti-correlated to increased astrocytes activation in GPe. Reduction of dopamine in GPe increases GABA tone through astrocytic D2-dependent mechanisms, resulting in inhibition of GPe neurons ^61^. This might re-establish the activity of SNc/VTA dopamine neurons, which are monosynaptically inhibited by GPe projecting neurons ^62^. Dopamine levels in basal ganglia are more closely related to motivational behavior than cell spiking of dopaminergic neurons in SNc/VTA ^63^. Nevertheless, astrocytes in SNc/VTA can also modulate the excitatory states of dopamine neurons ^48^, and the SNc, which is the primary source of dopamine in GPe, displays increased susceptibility to astrocytic D2 manipulations compared to VTA ^9^. In agreement, we demonstrated that deletion of astrocytic D2 either in GPe or SNc/VTA increased motivational behavior and increased dopamine levels in GPe. Thus, our findings well fit in this astrocytic-dopaminergic SNc-GPe homeostatic loop. Notably, in pathological conditions of astrocytic D2 overexpression (Dys1AGlast-/-, Fig. 7j), the motivation-related abnormal reduction of astrocytes activity in GPe would prevent the inhibition of local dopamine release and prevent the inhibition of GPe inhibitory projecting neurons. The loss of disinhibition of SNc/VTA dopamine neurons would prolong and delay dopamine dynamics in GPe, thereby leading to reduced motivated behavior (all of which was observed in our mice). Overall, our results extend previous evidence ^5, 10, 44^ by demonstrating that astrocytes are involved in the maintenance of motivational states and related ambient dopaminergic functionality in basal ganglia.

Increased astrocytic D2 in Dys1AGlast knockout mice was associated with loss of D2-dependent and delayed reward expectancy control of dopamine outflow. In contrast, neuronal overexpression of D2 enhances D2-dependent dopaminergic and behavioral responses ^64^. Moreover, we found that astrocytic D2 disruption increased basal ganglia dopamine outflow. In contrast, neuronal D2 disruption decreases striatal dopamine outflow ^65, 66^. Finally, in net contrast to astrocytic Dys1A deletion (our data), selective disruption of Dys1 in SNc/VTA dopaminergic neurons ^67^ resulted in altered locomotion, altered amphetamine-dependent dopamine release, and unaltered PPI. Thus, our findings reveal distinct astrocytic- *versus* neuronal-mediated control of dopamine signaling, at least within basal ganglia circuits.

In summary, we uncovered a selective, pivotal role for astrocytic Dys1A/D2 in the modulation of dopamine pathways within basal ganglia with direct implication for motivational and sensorimotor gating abilities relevant to schizophrenia. These findings provide a deeper mechanistic understanding of astrocytic regulation of dopamine signaling and its potential involvement in behavioral dysfunctions in psychiatric disorders. Indeed, our findings are of both fundamental and clinical relevance as they refine the concept of dopamine dysfunction in behavioral disorders and point to basal ganglia astrocytes as a promising target.

## Acknowledgments

We thank M. Morini, E. Albanesi, D. Cantatore, G. Pruzzo, T. Catelani, D. Mauro, S. Monari, F. Torri, B. Chiarenza, A. Monteforte and C. Chiabrera, for technical support. We thank GlaxoSmithKline and Dr. S. Wilson for generously donating the newly generated Dys1Aflox mice. We thank Dr. D.R. Weinberger and Dr. R.E. Straub for initial access to the Brain Cloud databank. We thank Dr. L. Tian and Dr. T. Patriarchi for generously sharing the initial dLight virus samples. This work was supported by funding from the Marie Sklodowska-Curie Fellowships (grant n.796244) to CD; the Ministero dell’Universita’ e della Ricerca italiano (project PRIN 2017K2NEF4) to FD; the Istituto Italiano di Tecnologia, the Brain and Behavior Research Foundation (2015 NARSAD grant n.23234), the Ministero della Salute italiano (project GR-2016-02362413), and Fondazione Telethon Italia (project GGP19103) to FP.

## Author Contributions

Conceptualization, RM, GT, FM and FP; Methodology and Investigation, RM, GT, DD, CD, VF, GO, RM, FM, GP, AF, MADL, FM and FP; Resources, GO, DR, JLW, GML, CSW, FM and FP; Writing, all authors; Visualization, RM, GT, DD, CD, GL, GO, RM, FM and FP; Supervision, FM and FP; Funding Acquisition, FD, CSW, and FP.

## Online Methods

### Mice

All procedures were approved by the Italian Ministry of Health (permit n. 230/2009-B, 107/2015-PR, and 749/2017-PR) and local Animal Use Committee and were conducted in accordance with the Guide for the Care and Use of Laboratory Animals of the National Institutes of Health and the European Community Council Directives. Routine veterinary care and animal maintenance were provided by dedicated and trained personnel. Male and female littermate mice between 3-7 months of age were used throughout. Animals were housed 2-4 per cage, in a climate-controlled animal facility (22°C ± 2°C) and maintained on a 12-hr light/dark cycle (08:00 on; 20:00 off), with food and water available *ad libitum.* The experimenter handled the mice on alternate days during the week preceding the tests. Body weights and general appearance of mice were recorded before starting behavioral testing.

#### Dys1, Dys1A, Dys1AGlast and D2 mutant mice

The Dys1 heterozygous mutant mice (Dys1+/−) and their wild-type littermates (Dys1+/+) with a C57BL6/J background were bred and used as previously described ^20, 26^. The Dys1A_flox/flox_ mice generated by Glaxo SmithKline ^27^ were kept on C57BL6/J background and presented two loxP sites flanking exon 5, which is necessary for correct expression of the Dys1A long isoform. Constitutive Dys1A deletion (Dys1A-/-) or partial reduction (Dys1A+/-) was obtained crossing Dys1A_flox/flox_ mice with a germline Cre deleter transgenic strain (Taconic-Artemis Germany). The breeding scheme used consisted of mating one male Dys1A+/- with two Dys1A+/- females. Astrocyte-specific loss (Dys1AGlast-/-) or partial reduction (Dys1AGlast+/-) of the expression of Dys1A was obtained by mating Dys1A_flox/flox_ mice with the inducible Cre transgenic line Glast-CreERt2 (Mori et al. 2006). The resulting strain was then crossed with the reporter line stop_flox/flox_ tdTomato (Jackson Laboratory). Tamoxifen (T-5648, Sigma-Aldrich, St. Louis, MO, USA) was dissolved in corn oil (Sigma-Aldrich, C-8267) and 1:20 ethanol at 55°C for two hours to prepare a stock solution at 100mg/ml. Stock solutions were aliquoted and stored at -20°C. To induce Cre-mediated recombination, mice aged 60 days were administered tamoxifen 5mg/day for two consecutive days by oral gavage (Mori et al. 2006). Testing started 30 days later. The breeding scheme consisted of mating one Dys1A_flox/+_ Glast-CreERt2_tg/tg_ tdTomato_flox/flox_ male mouse with two Dys1A_flox/+_ Glast-CreERt2_+/+_ tdTomato_flox/flox_ female mice. Genotyping was performed by PCR using wild-type, targeted and Cre allele-specific primers (Tab.1). Drd2^loxP/loxP^ mice were purchased from Jackson Laboratory (stock #020631)^68^, and the colony was maintained in homozygosity. All mutant mice used were viable, fertile, normal in size and did not display any gross physical or behavioral abnormalities.

### Behavior

#### Locomotor Activity

Mice were tested in an experimental apparatus consisting of four gray, opaque open field boxes (40×40×40 cm) evenly illuminated by overhead lighting (5±1 lux). Each session was video-recorded using an overhead camera from ANY-maze (Stoelting Co., Wood Dale, IL, USA) with the experimenter absent from the room during the test. Activity was tracked during the first exposure to the empty open field arena for 30 minutes. For amphetamine experiments, mice were tested in the same open field arenas. First, mice were placed in the empty open field and allowed to explore for 10 minutes. Then, mice were removed from the arena, injected with 1.5 mg/kg/10ml D- amphetamine sulphate (i.p.; Sigma-Aldrich) and placed back in the open field for an additional 60 minutes. This procedure was repeated for five consecutive days. All sessions were videotaped and tracked with ANY-maze software (Stoelting Co.).

#### Male-Female social interaction

The test was conducted in Tecniplast cages (35×23×19 cm) illuminated (5±1 lux) and video-recorded using a Unibrain Fire-i digital camera. The video camera was mounted facing the front of the cage to record the session for subsequent scoring of social investigation parameters as previously described ^69^. Unfamiliar female stimulus mice in estrus were matched to the subject male mice by age and maintained in social groups of four per home cage.

#### Social habituation/dishabituation task

Naıve mice were tested in Tecniplast cages (35×23×19 cm) illuminated (5±1 lux) and video-recorded using a Unibrain Fire-i digital camera. As described previously ^69^, mice were placed individually for environmental habituation to the test cage 1h prior to testing. A stimulus mouse (unfamiliar of the same sex) was introduced into the testing cage for a 1- min interaction. At the end of the 1-min trial, we removed the stimulus animal and returned it to an individual holding cage for 3 minutes. We repeated this sequence for three trials with 3-min intertrial intervals. In a fifth ‘dishabituation’ trial, we introduced a new (unfamiliar) stimulus mouse into the testing cage. Videos of behaviors were recorded and subsequently scored offline.

#### Attentional Set-Shifting Task

Attentional set-shifting was tested in the two-chamber ID/ED Operon task as previously described ^20, 32^. After random selection of mice for the ID/ED task, all behavioral parameters were obtained blind to the genotype of the animals. For habituation to the apparatus, during the first 2 days, mice were habituated for 45 min to the apparatus with only neutral stimuli (Habituation 1) and trained to move from one chamber to the other (Habituation 2). Any nose poke into the nose-poke holes resulted in a pellet delivery into the food receptacle. The next day, mice were trained to perform two randomly presented simple discriminations (e.g. between smooth vs. sand cardboard; light on vs. light off; peach vs. sage) so that they were familiar with the stimulus dimensions (Habituation 3). These exemplars were not used again. The mice had to reach a criterion of eight correct choices out of ten consecutive trials to complete this and each following testing stage. Performance was measured in all phases of all experiments using number of trials to reach the criterion; time (in minutes) to reach the criterion and time (in seconds) from breaking the photobeams adjacent to the automated door to a nose-poke response (latency to respond). A session started when a mouse was placed in one of the two chambers where all the stimuli were neutral. Then the transparent door was dropped to give the mouse access to the other chamber where the stimuli cues were on. The series of stages comprised a simple discrimination (SD), compound discrimination (CD), compound discrimination reversal (CDRe), IDS, IDS reversal (IDSRe), a second IDS (IDS2), IDS2 reversal (IDS2Re), EDS, and EDS reversal (EDSRe). The mice were exposed to the tasks in this order so that they could develop a set, or bias, toward discriminating between the correct and incorrect nose poke hole.

#### Acoustic startle response and prepulse inhibition (PPI)

Acoustic startle response and PPI were measured using SR-Lab Systems (San Diego Instruments, San Diego, CA, USA) and TSE Startle Response System (TSE Systems GmbH, Bad Homburg, Germany) following previously described protocols ^26, 70^. Briefly, a sudden acoustic stimulus (120 dB) elicits the startle response, while an acoustic, non-startling pre-pulse (74; 78; 82; 86; 90 dB) preceding the startle stimulus inhibits the startle response (PPI). The startle response elicited by sudden sensory stimuli and its PPI are among some of the most widely studied phenotypes that are highly conserved across mammalian species. A background level of 70 dB white noise was maintained throughout the test session.

#### Progressive Ratio Test

We tested mice in a motivational nose-poke operant paradigm for 14 mg 5- TUL pellets (Test Diet) as described previously ^41^. To avoid confounding factors linked to food restriction/deprivation experience, mice were always provided with food and water *ad libitum*. The operant chambers used (MED Associates Inc, VT, USA) were equipped with two nose-poke holes mounted at the left and right of a central food magazine, each equipped with infrared photobeams connected to a computer with MED-PC V software. Nose poking into one of the two holes resulted in pellet delivery (active hole), whereas nose poking into the other hole (inactive hole) triggered the house light for 5 seconds (left and right randomly assigned and balanced between groups). Free water was available all time via a water bottle dispenser. Pellets were delivered to the food magazine by an automated dispenser situated outside the experimental chamber. Mice were placed into the operant chambers in the evening around 5pm and taken out the following morning around 9-10am. Lights within the sound attenuating boxes in which the operant chambers were located ensured mice experienced a light/dark cycle the same as that of holding rooms. Training and testing started automatically from the beginning to the end of the dark phase (8pm to 8am). Initially, a fixed ratio (FR)-1 reinforcement schedule was applied, i.e. one nose poke in the active hole resulted in delivery of one pellet. Mice were exposed to the FR1 schedule until they reached the criterion of > 80% active pokes during the entire night for two consecutive nights.

Mice that met this learning criterion were switched to a FR3 reinforcement schedule, i.e. three nose-pokes in the active hole produced delivery of one pellet. The FR3 reinforcement schedule lasted two nights if mice met the criterion of > 80% active pokes during the entire night. Afterward, mice were exposed to a progressive ratio (PR) schedule that lasted two hours from the beginning of the night phase and was changed nightly; first night: PR3; second night: PR6; and third night: PR9. Mice were returned to their home cage after the PR test. During the PR experiment, the number of active nose-pokes required to obtain each successive food pellet was progressively increased by three (PR3, 3n+3), six (PR6, 6n+6) and 9 (PR9, 9n+9; where n=number of pellets earned). For example, in PR3 earning the first reinforcer required three active nose pokes, the second six nose pokes, the third nine nose pokes, etc. Likewise, in PR9 earning the first reinforcer required 9 active nose-pokes, the second eighteen nose-pokes, the third twenty-seven nose-pokes, etc. Following each PR session, we calculated the breakpoint (BP) as the last ratio level completed before the end of the two-hour testing session. For example, under the PR3 or the PR9 reinforcement schedules, to earn the third food pellet a mouse had to poke 3+6+9 or 9+18+27 times in the active hole, and thus was given a BP value of 9 or 27, respectively. The BP is a well-validated measure reflecting the strength of the reinforcer and the motivational state of the animal.

### Immunohistochemistry

Mice were deeply anesthetized (urethane 20%) and perfused transcardially with PBS followed by 4% formaldehyde solution (Sigma-Aldrich) in PBS, pH 7.4. Brains were extracted, post fixed overnight in 4% formaldehyde and cryoprotected in 30% sucrose in PBS. 40-μm-thick coronal sections containing region of interest were cut on a freezing microtome (VT1000S, Leica Camera AG, Wetzlar, Germany) and collected in PBS before being processed for immunohistochemistry. For GFAP immunostaining, free-floating slices were first washed once in 0.3% Triton X-100 PBS (PBS-T) for 10 minutes, and twice with 0.1% PBS-T, then incubated for 1 h in a blocking solution of 5% normal goat serum (NGS) in 0.1 % PBS-T. Subsequently they were incubated overnight at 4 °C with 1:300 rabbit polyclonal anti-GFAP antibody (Novus Biologicals, Centennial, CO, USA) in blocking solution. For TH immunostaining, slices were blocked in PBS with 2% bovine serum albumin (BSA) and 0.2% Tween-20 (Sigma-Aldrich) for 1 hour at room temperature, and then incubated with 1:1000 rabbit anti-TH primary antibody (AB152, Millipore, Burlington, MA, USA) in blocking solution for 72 hours at 4°C. Slices were then incubated with Goat anti-rabbit Alexa 488 secondary antibody, 1:1000 for GFAP stained slices, and 1:500 for TH stained slices (A-11035, Invitrogen by Thermo Fisher Scientific, Waltham, Massachusetts, USA) for 1 hour at room temperature. Between steps four washes in blocking solution were applied for 10 minutes. Slices were then mounted with ProLong™ Gold Antifade Mountant (ThermoFisher Scientific) and imaged in an inverted laser scanning confocal microscope (A1 Nikon, Shinjuku, Japan) using a 20x or 40x objective. Quantification and analysis were performed using Fiji software (Wayne Rasband, NIH, USA), outlining regions of interest. To analyze the number of Glast-positive cells in GPe sections, 20x magnification confocal images were acquired. For each animal three images were taken from sliced collected between 0.58 and -0.70 mm from Bregma. Glast-positive cells were quantified by averaging the cell density within and across each animal from 10µm maximum projections images. Counts were performed using 3D objects counters tool provided by NIH Imagej software and the subsequent analyses were performed following a blind procedure. Astrocyte surface and relative intensity were calculated analyzing GFAP signal in GFAP- and Glast-positive cells from 10µm maximum projection images, acquired at 40x magnification.

### Electron microscopy

Mouse brains were perfused in 4% formaldehyde and 2% glutaraldehyde and embedded as described previously (Polishchuk & Mironov 2001). Bright field transmission electron microscopy images were acquired from thin (70 nm) sections using a Gatan Orius SC1000 series CCD camera (4008 x 2672 active pixels) (Gatan, Pleasanton, USA), fiber optically coupled to high-resolution phosphor scintillator under a JEOL JEM-1011 transmission electron microscope (TEM) (JEOL, Tokyo, JAPAN) with thermionic source (W filament) and maximum acceleration voltage 100 kV. All transverse sections of the Golgi Complex (GC) were taken at the same magnification (X6000) and analyzed using point-counting procedures, with surface densities of Golgi Complex (Sv_i_GC) and Cytoplasm (Sv_i_CYT) determined according to Leitz ASM system. Moreover, a qualitative score from 1 to 3 was assigned to all GC: the maximum score (3) was given when finding a group of cisternae organized in stacks containing tubular and vesicular structures, as defined for GC, and the lower score (1) was given when GC structure was destroyed. Double tilt high angular annular dark field (HAADF) scanning TEM (STEM) tomography was performed using a Tecnai F20 transmission electron microscope (FEI Company, Eindhoven, The Netherlands), equipped with a field-emission gun operating at 200 kV and a Gatan Ultrascan US1000 (Gatan, Pleasanton, USA) . For the reconstruction of the Golgi apparatus in Dys1+/- mouse astrocytes a 300-nm-thick section was tilted over ±60° with the following tilt scheme: 1° at tilt higher than ±30° and 2° intervals at intermediate tilts. The images were acquired using a HAADF detector at a magnification of 40,000 times. Computation of double tilt tomogram was done by combining two tilt series taken around two orthogonal axes with the IMOD software package. 3D reconstruction has been performed using Amira™ Software (Thermo Fisher Scientific).

### Cell cultures

Astrocytes-enriched cell cultures were obtained from cortices dissected from PND1 mice. Pups were sacrificed by cervical dislocation and cortices were quickly dissected in ice-cold HBSS (Hanks’ Balanced Salt Solution, Gibco ThermoFisher Scientific). Samples were incubated in HBSS with 0,125% Trypsin-EDTA (ThermoFisher Scientific) and 1 mg/mL DNAase I (Sigma-Aldrich) for 20 minutes at 37°C. A solution of DMEM (Dulbecco’s Modified Eagle Medium, Gibco ThermoFisher Scientific) with 10% horse serum and 1% Penicillin-Streptomycin (Sigma-Aldrich) was added to the samples, which were then centrifuged 1200 rpm and washed twice in complete medium. Samples were dissociated mechanically in complete medium and filtered through 40 μm cell strainers. Cell suspensions were finally plated on poly-D-lysine coated plates. Cells were cultured until 100% confluence. Neuronal cell cultures were obtained from E18 mice embryos. Cortices were dissected in ice-cold HBSS and incubated in HBSS with 0,125% Trypsin-EDTA and 0,25 mg/mL DNAase I for 30 minutes at 37 °C. A solution of Neurobasal^TM^ medium (Gibco ThermoFisher Scientific) with 10% inactivated fetal bovine serum (Signa-Aldrich), 1% Penicillin-Streptomycin, 1% GlutaMAX™ Supplement and 2% B27™ Supplement (Gibco ThermoFisher Scientific) was added to the samples, which were then centrifuged 1200 rpm and resuspended in complete medium before mechanical dissociation. Samples were filtered with 40 μm cell strainers, centrifuged 700 rpm and resuspended in complete medium. Cell suspensions were finally plated on poly-D-lysine coated plates. Neurons were cultured until complete maturation.

### Western Blot

For western blot analysis of Dys1 isoforms, we used Dys1+/+ mice at embryonic stage E14.5, postnatal day 7, 35 and 90, Dys1 and Dys1A knockout mice, glial and neuronal cell cultures. Animals were sacrificed by cervical dislocation; brains were rapidly dissected and stored at -80°. Tissues or cultured cells were lysed in RIPA buffer and Protease Inhibitor Cocktail (Sigma-Aldrich). Western blot analysis was performed using mouse polyclonal anti-Dysbindin antibody (PA311 validated and produced by ^15^ and the anti-actin antibody (Sigma Aldrich). Amounts of 25μg of protein from precipitated homogenates were separated on SDS-PAGE, electrotransferred onto nitrocellulose membranes, and then probed with primary antibodies: mouse monoclonal anti-Dysbindin antibody (1:1000) and mouse anti-Actin antibody (1:10000). Immune complexes were detected using appropriate peroxidase-conjugated secondary antibodies (Thermo Fisher Scientific) and a chemiluminescent reagent (ECL prime; GE Healthcare Europe GmbH, Milan, Italy). Densitometric analysis was performed using ImageQuantTL software (GE Healthcare Europe GmbH). Results were normalized to respective control conditions.

### Slices surface biotinylation

These experiments were performed as previously described ^20, 26^. Mice were anesthetized with isoflurane and decapitated. The brain was sectioned in cold carb-oxygenated HBSS enriched with 4mM MgCl, 0,7 mM CaCl_2_ and 10 mM D-glucose, equilibrated with 95% O2 and 5% CO2 to yield a pH 7.4) on a vibrating microtome at a thickness of 300 μm. Striatum and prefrontal cortex were dissected from coronal slices. Before starting the surface biotinylation reaction, and to ensure a gradual cooling of the cells, the tissues were washed twice for 5 min in ice-cold HBSS buffer. The filters holding the tissues were transferred to a well containing an excess of biotinylation reagent solution of 100 μM NHS-LC-biotin (Pierce, Appleton, WI, USA) in HBSS. After 45 min of incubation, the tissues were transferred to another well and washed twice with the HBSS buffer containing 200 mM Lysine (Sigma-Aldrich), to block all reactive NHS-LC-biotin in excess. The tissues were washed twice with ice-cold HBSS and immediately placed on ice to mechanically disrupt the tissue in 120 µl of lysis buffer (1% TX-100, PBS1X and a cocktail of protease inhibitors (Sigma-Aldrich). To discard extra debris, homogenates were centrifuged for 5 min at 4°C at 13.000 r.p.m. and supernatants were collected. To precipitate the biotinylated proteins from the homogenates 50 µl of immobilized Streptavidin beads (Pierce) were added to the samples and the mixture was rotating for three hours at 4°C. The precipitates were collected by brief centrifugation, mixed with 50 µl of SDS-PAGE loading buffer, boiled for 5 minutes and stored at -80°C until use. Protein extracts were separated on precast 10% SDS/PAGE (Biorad, Milan, Italy) and transferred to nitrocellulose membranes. Blots were incubated with primary antibodies overnight at 4°C. Antibody used were dopamine D2 receptor (sc- 5303, Santa Cruz Biotechnology, Dallas, TX, USA) and (AB5084P, Millipore), Synaptophysin (sc- 365488, Santa Cruz Biotechnology) and Transferrin Receptor (sc-21011, Santa Cruz Biotechnology). Immune complexes were detected using appropriate peroxidase-conjugated secondary antibodies (Thermo Fisher Scientific) and a chemiluminescent reagent (ECL prime; GE Healthcare Europe GmbH, Milan, Italy). Densitometric analysis was performed by ImageQuantTL software (GE Healthcare Europe GmbH). Results were normalized to respective control conditions.

### Stereotaxic injections

All surgeries were performed under aseptic conditions. Mice were deeply anesthetized with a mix of isoflurane/oxygen (2%/1%) by inhalation and mounted into a stereotaxic frame (David Kopf Instruments, Tujunga, CA, USA). Following shaving and preparation of the skin, a cranial hole was made above the targeted area. All measurements were made relative to bregma, in accordance with the mouse brain atlas ^71^. The viral injection was performed using a borosilicate pipette at a rate of 50nl/min using a 10-μL Hamilton syringe. After each injection, 10-15 minutes were allowed before slowly withdrawing the micropipette. Animal receiving dLight injection (AAV-CAG-dLight1.2, 400nl) in the GPe (Coordinates in mm: virus: AP: -0.4, ML: 1.9 and DV: 3.7, optic fiber implant: AP: 0.4 mm; ML: ± 1.9 mm; DV: -3.6 mm) were implanted with a custom-made optic fiber (200µm, 0.50NA, Thorlabs, Newton, NJ, USA) during a second surgery.

For cre expression in astrocytes, AAV viral injections (AAV8-GFAP-Cre-GFP or AAV8-GFAP- GFP) targeted the SNc/VTA (100nl, AP: -3 mm; ML: ± 0.50 mm; DV: -4.7 mm) or the GPe (400nl, AP: -0.4, ML: 1.9 and DV: 3.7). For calcium imaging viral AAV injections (AAV5.GfaABC1DcytoGCaMP6f.SV40 or GfaABC1D.cyto-tdTomato.SV40, Addgene, Watertown, MA, USA) targeted the PFC (300nl, AP: +1.9 mm; ML: ±0.3 mm; DV: -2.4 mm), the GPe (2×400nl, AP: -0.4, ML: 1.9 and DV: 3.7) or the SNc/VTA (250nl, AP: -3.1 mm; ML: ± 0.75 mm; DV: -4.5 mm), together with the control virus AAV-GFAP-TdTomato (1:5 ratio from the GCamP volume).

### Chromatographic analyses of dopamine, DOPAC, HVA, NA, 5HT, and 5HIAA

#### Ex vivo tissue collection

Brains were harvested following rapid decapitation and sliced in 1 mm sections in a chilled stainless-steel mouse brain matrix. Slices were frozen on glass slides mounted on dry ice. Using a 2 mm biopsy punch, bilateral PFC, STR and GPe punches were collected accordingly to the mouse brain atlas ^71^, and stored at -80°C until neurochemical analyses.

#### In vivo microdialysis

Microdialysis procedure was performed as previously described ^20, 26^. A concentric dialysis probe with a dialyzing portion of 1 mm was prepared and stereotaxically implanted in the right GPe (coordinates of the dialyzing portion tip, in mm, relative to the bregma point, according to the atlas of Paxinos and Watson, 2001: anteroposterior (AP)=-0.4, lateral (L)=+1.9, ventral (V)=-4.5) under isoflurane anesthesia. After surgery, mice were housed individually to recover for 24 hours. On the day of microdialysis, probes were perfused at a constant flow rate (1 μl/min) by means of a microperfusion pump, with artificial cerebrospinal Fluid (aCSF, in mM: 147 NaCl, 4 KCl, 2.2 CaCl_2_). After 30 min stabilization, samples were collected every 20 min and stored in dry ice until the end of the experiment. Three groups of three dialysates (one hour per group) were consecutively collected: “baseline” (aCSF perfusion), “quinpirole” (perfusion of 25 nM quinpirole), and “wash-out” (aCSF perfusion). At the end of the microdialysis experiment, brains were collected and sliced to check probe implantation (only data obtained from mice with probes correctly implanted in the GPe were included in the results).

#### Quantification of monoamines and metabolites by HPLC

PFC, STR, and GPe tissue samples were lysed by sonication in 0.1 M perchloric acid, and centrifuged (15,000 x g, 10 minutes, 4°C). The supernatant was filtered by centrifugation (20,000 x g, 5 min, 4°C) in ultra-free microcentrifuge tubes (Millipore, Burlington, Massachusetts, USA). Supernatants obtained from PFC, STR or GPe samples, and from dialysates obtained from GPe *in vivo* microdialysis were injected (11 μl) into a high-performance liquid chromatography apparatus (Alexys UHPLC/ECD Neurotransmitter Analyzer, Antec Scientific, Zoeterwoude, The Netherlands), equipped with an autosampler (AS 100 UHPLC, micro, 6-PV, Antec Scientific). The mobile phase [containing (in mM) 100 phosphoric acid, 100 citric acid, 0.1 EDTA.2H 2H_2_O, 3 octanesulfonic acid. NaCl plus 8% acetonitrile, adjusted to pH 3.0 with NaOH solution (50%)] was delivered at 0.050 ml/min flow rate with a LC 110S pump (Antec Scientific) through an Acquity UPLC HSS T3 column (1 x 100 mm, particle size 1.8 µm; Waters, Milford, Massachusetts, USA). Detection of dopamine, DOPAC and HVA was confirmed and carried out with two system. An electrochemical detector (DECADE II, Antec Scientific) equipped with a Sencell with a 2 mm glassy carbon working electrode (Antec Scientific) set at +600 mV versus Ag/AgCl. Output signals were recorded with Clarity (Antec Scientific). The second HPLC was equipped with a reversed-phase column (C8 3.5 um, Waters, USA) and a coulometric detector (ESA Coulochem III; Agilent Software). The electrodes of the analytical cell were set at +350 mV (oxidation) and −200 mV (reduction). The mobile phase contained 50 mM CH_3_COONa, 0.07 mM Na_2_EDTA, 0.5 mM noctyl sulfate, and 12% (v/v) methanol, the pH of mobile phase was adjusted with CH_3_COOH to 4.21. The sensitivity of the assay for DA/DOPAC/HVA was 5 fmol/sample. For tissue sample analysis, data were normalized by tissue weight. Dialysate contents were converted into percentages of the average baseline level calculated from the three fractions of the first hour of collection (“baseline” period), and are expressed as averaged percentages of “baseline”, “quinpirole” and “wash-out” periods, obtained in each experimental group.

### Fluorescence-activated cell sorting and cytometry

#### Sorting of tdTomato-positive/negative cells

Dys1AGlast mice were sacrificed by decapitation 4 weeks post Tamoxifen administration. Brains were quickly removed from the skull, and sub-cortical regions containing basal ganglia were immediately dissected and washed in ice-cold PBS with 5.5 mM D-Glucose and 0.32 mM Sodium Pyruvate. Samples were then processed with the Adult Brain Dissociation Kit and MACS^®^ SmartStrainers 70 µm diameter cell strainers (Miltenyi Biotec, Bergisch Gladbach, Germany) to be disaggregated, filtered, and removed from myelin, debris and red blood cells, according to manufacturer’s instructions. Final single-cell suspensions were obtained in 200 µL PBS with 0.5% BSA and 25 mM HEPES (sort buffer); to remove remaining debris and aggregates, each cell suspension was filtered through a CellTrics^®^ 50 µm mesh diameter cell strainer (Sysmex, Norderstedt, Germany). Samples were analyzed with BD FACSAria^™^II cytometer and cell sorter, by using FacsDIVA software (BD Biosciences, Franklin Lakes, New Jersey, USA). TdTomato-positive and -negative astrocytes were gated on singlets and separately collected into 20 µL sort buffer. Non-fluorescent cells were used as negative control for gating.

#### Cytometry for D2 expression quantification

Cell suspensions were obtained as described in the previous paragraph. Cells were stained with 0.25 µL of eBioscience™ eFluor™ 780 viability dye (Invitrogen by Thermo Fisher Scientific) for 30 minutes at 4°C in the dark and washed in 2 mL PBS with 0.5% BSA. Cells were then resuspended and incubated for 5 min in PBS with 2% BSA and stained with 2 µg Anti-D2 Dopamine Receptor-FITC Antibody solution (Alomone Labs, Jerusalem, Israel) for 40 minutes, at 4°C in the dark. Finally, cells were washed in 2 mL PBS with 0.5% BSA and resuspended in 250 uL sort buffer. Samples were analyzed with BD FACSAria^™^II cytometer and cell sorter, by using FacsDIVA software (BD Biosciences). FITC mean fluorescence intensity (MFI) was measured on tdTomato-positive and negative cells, which were previously gated on 20000 living cells.

### Quantitative real-time PCR

Sorted astrocytes were pelleted by centrifugation at 1000 rpm for 5 minutes at 4° C, and total RNA was extracted using the RNeasy MicroKit (QIAGEN, Hilden, Germany), according to manufacturer’s instructions. RNA quantification and quality assessment were conducted with Nanodrop ND1000 microspectrophotometer (Thermo Fisher Scientific). RNA was retrotranscripted in cDNA using the High-capacity cDNA Reverse Transcription Kit (Applied Biosystems, Foster City, California, USA), according to manufacturer’s instructions. Relative gene expression was assessed by real-time qPCR (7900 HT, Applied Biosystems), and 96-well plates were used for amplification reactions in 10 µL final volume per well, charged as follows: 5 µL iTaq™ Universal SYBR® Green Supermix (Bio-Rad, Hercules, California, USA), 1 µL primers mix and 4 µL template, containing 2-5 ng cDNA; primers were used at 0.5 µM concentration. The thermal profile used was the following: 30 sec at 95°C for denaturation, 15 s at 95° C and 1 min at 60 °C (40 cycles) for amplification, and 15 s at 95 °C, 15 s at 60 °C and 15 s at 95 °C for dissociation. All reactions were performed in triplicate and analyzed by SDS 2.3 software (Bio-Rad) to calculate cycle threshold (Ct) values. Relative gene expression levels were normalized with Gapdh as reference gene and compared between genotypes with the ΔΔCt method. The following primers were used: Glast forward 5’-CTCACGGTCACTGCTGTCAT- 3’ reverse 5’-GCCATTCCTGTGACGAGACT-3’; Dys1A forward 5’-ATGGCAA- GCCTGGCTCATTTAGA-3’ reverse 5’-AGTCCTCCAGGTGCAGCAAAT-3’; NeuN forward 5’- ATCGTAGAGGGACGGAAAATTGA-3’ reverse 5’-GTTCCCAGGCTTCTTATTGGTC-3’; D1 forward 5’-GACATACGCCATTTCATCCTCC-3’ reverse 5’-ATGCGCCGGATTTGCTTCT-3’; D2 forward 5’-ACCTGTCCTGGTACGATGATG-3’ reverse 5’-GCATGGCATAGTAGTTGTAG- TGG-3’; D3 forward 5’-TGGGGCAGAAAACTCCACTG-3’ reverse 5’-TACCAGACCGTT- GCCAAAGAT-3’; Dat forward 5’-AAATGCTCCGTGGGACCAATG-3’ reverse 5’- GTCTCCCGCTCTTGAACCTC-3’; Vmat2 forward 5’-ATGCTGCTCACCGTCGTAG-3’ reverse 5’-GGCAGTCTGGATTTCCGTAGT-3’; Maob forward 5’-ATGAGCAACAAAAGCGATGTGA-3’ reverse 5’-TCCTAATTGTGTAAGTCCTGCCT-3’; Oct3 forward 5’-CAGCCCGAC- TACTATTGGTGT-3’ reverse 5’-TGAGCTGGTATTAGTGGCTTCC-3’; Th forward 5’- GTCTCAGAGCAGGATACCAAGC-3’ reverse 5’-CTCTCCTCGAATACCACAGCC-3’; Gapdh forward 5’-GTGATGGGTGTGAACCACGA-3’ reverse 5’-CTGTGGTCATGAGCCCTTCC-3’.

### Fiber photometry in behaving mice

#### In vivo dopamine dynamics in freely moving mice

Seven days after the initial viral injection of AAV- dLight, a flat-cut multimode optic-fiber (200µm, 0.50NA, Thorlabs) was gently inserted 300-400µm above the virus injection site in the GPe. The implant was maintained in position using 2-to-3 anchor screws and affixed to the skull using dental cement, a second ferrule was implanted contralaterally as a dummy cannula to reduce rotation and movement artifacts.

#### In vivo calcium imaging in freely moving mice

Seven to 10 days following the injection of AAV5.GfaABC1DcytoGCaMP6f.SV40 and AAV-GFAP-TdTomato, mice underwent a second surgery for the implantation of two flat-cut multimode optic-fiber (200µm, 0.50NA, Thorlabs) 300- 400µm above the virus injection site of the GPe or the VTA. The implant was maintained in position using 2-to-3 anchor screws and affixed to the skull using dental cement.

#### Fiber photometry recordings

Seven days after the surgery, mice were trained in the progressive ratio test as described above. The day following two consecutives overnight FR3 sessions with more than 80% of correct choices, mice were transferred to a 1-hour day session at FR3 schedule, until completion of 2 consecutive days at more than 80% correct choices. At that stage, all mice were connected to the patch-cord, with no recording to habituate the mice. Next, mice were tested at the same conditions in consecutive daily sessions for FR3, PR3, PR6 and PR9 schedules, and this time calcium signal was acquired. The fiber photometry signal was acquired for 10 minutes before the beginning of the sessions to minimize photobleaching effects. tdTomato fluorescence in astrocytes was used as control signal. Two LED beams (465 and 560nm, Doric Lenses, Quebec, Canada) were focused onto an integrated fluorescence mini cube (for dLight #Ifmc5, Doric lenses) or two fluorescence mini cube (for GCamP, #Ifmc5, Doric lenses) using a 50:50 beam divider (Thorlabs) where both excitation wavelengths were filtered at 460-490 and 555-570nm respectively. Excitation wavelength were adjusted to reach approximately 30µW to each ferrule with variation between animals based on the signal-to-noise-ratio (SNR). To measure F_0_, both beams were pulsed at anti-synchronous frequency (330Hz for 560nm, and 210Hz for 465nm). The beams were redirected into a low-autofluorescence 400µm patch-cord and connected to the head-implant using a ceramic sleeve. For dLight experiments, a dummy patch-cord was connected to the dummy cannula, limiting rotation and movement artifacts. The resulting emission signals were merged and filtered at 500-540nm (GCAMP) and 580-680nm (tdTomato). For single site photometry (dLight) both signals were directed onto their own femtowatt photosensor (Newport, Irvine, CA, USA) and deconvolved directly using their respective excitation wavelength. For dual site photometry (GCamP) the resulting signal of each ferrule were directed onto a single femtowatt photosensor and analyses using lock-in deconvolution. Low-DC signal of the photosensors were then processed using RZP5 hardware and Signal software (Synapses, Tucker Davis Technology, Alachua, FL, USA). The acquisition rate was down sampled at 1017Hz (6^th^ order) and both signals (dLight/tdTomato or GCAMP/tdTomato) were first deconvolved using the excitation wavelength with the time period of laser OFF (F0) and laser ON (ΔF). Behavior events output from the operant boxes (Med Associates) were duplicated, converted to TTL signal (5V) using a custom-made 28-to-5V converter and sent to the inputs port of the acquisition box. Behavioral and florescent signal extracted from the Synapse software were then converted to Matlab using TDTtoolbox.

#### Analyses of the photometry signal

All analyses were done on Matlab (2020a, Mathworks). For each animal, recordings were done during 1 hour daily sessions. Signal for the FR3 and PRs schedules were analyzed separately before to be merged, as presenting similar response patterns. We extracted the timestamps of each event using the TTL onset: entry in a correct poke, entry in the food magazine, and entry in the incorrect hole. Signal processing was done for each behavioral event aligning the GCamP/dLight and the tdTomato signal with a 120sec window centered at the behavioral event (1mins baseline, 1min post-event), and then grouped by trial type, genotype and condition. Each signal trials were then analyzed as follow: all signals were detrend using a linear detrend function and z-score using the pre-event windows as mean and standard-deviation value. All data point (F) were normalized to the 60s windows (F_0_) to express calcium variation (ΔF/F_0_) where ΔF is the change of calcium between F and F_0_. Using such an approach allow us to conserve the SNR and the subthreshold calcium oscillation while not being affected by inter-animal variation nor the low SNR of the astrocytic calcium signal. Following completion of the analyses, signal was down sampled to a 2Hz frequency. For statistical analyses, a non-overlapping moving window of 500ms were used to extract the individual trials signal prior and after each event. For each behavioral event, a repeated measure ANOVA was used to analyze variation of GCamP, dLight and tdTomato signals from the baseline and between genotypes. In order to avoid the extraction of non-specific signal, random trials for each animal were extracted using a 200 random iteration (timestamps) per animals and pooled as random lag for each event. All experiments were conducted blindly, and repeated in at least two different cohort of mice, analyzing at the same way the tdTomato, GCamP and dLight signals. We did not normalize GCamP/dLight signal to their tdTomato control signal due to the risk to increase non-specific signal (Joon Lee et al., 2019).

#### Signal correlation from photometry

For all behaviors, the correlation between GPe-signal and dLight-signal before and after magazine entries or correct pokes was counted based on the Pearson’s correlation coefficient (PCC). The overall activities of GCamP-GPe and dLight-GPe signal from 5 sec before to 5 sec after magazine entries or correct pokes were used to count overall PCC and their relative p-value. To understand the synchrony of the two signals after the magazine entries or correct pokes, the cross-correlation between GCamP-GPe and dLight-GPe was computed using a Matlab function (xcorr). This function calculates PCC between two signals when one of them is time shifted (lag) in respect to the other. It is possible to understand the time in which the traces are most correlated (lag) to each other by observing the PCC’s change linked to the time shift. If the highest PCC’s (absolute value) is displayed with no time shift (in lag 0), both signals are synchronized (within a window related to calcium oscillation decay and rising time).

### Drosophila

#### Stocks and crosses

The UAS-Ddysb-RNAi *Drosophila* line (v34355) used in this study was obtained from VDRC (Vienna Drosophila Stock Center). The Gal4 activator lines tubulin-Gal4 (5138), repoGal4 (7415), and elav-Gal4 (458), and the transgenic lines UAS-GalT-GFP (30902), were obtained from the Bloomington Stock Center, Indiana University. Experimental crosses were performed at 28°C.

#### Immunochemistry

*Drosophila* immunostaining was performed on wandering third instar larvae reared at 28°C. Third-instar larvae were dissected in PBS and fixed in 4% paraformaldehyde (PFA) in PBS for 15 minutes, washed in PBS 0,1% Triton X-100 (PBTX), and incubated with primary antibody overnight, and secondary antibody for 1 hour. The primary antibody anti-Repo-8D12 (1:200, DSHB) was used. Secondary antibody Cy5coniugated Goat anti-Mouse IgG (115-175-003) was from Jackson Immuno Research, and was used at 1:500. Third instar larvae were then mounted with Mowiol 488 and were imaged using a Nikon EZ-C1 confocal microscope equipped with a Nikon Plan APO 60.0×/1.40 oil immersion objective. Z-stacks with a step size of 1 µm were taken using identical settings. Each stack consisted of 15 to 20 plane images of 10 animals per genotype. The images obtained were processed and analyzed using *ImageJ*.

#### qRT-PCR

*Drosophila* samples (8 mg each) were homogenized in TRIzol Reagent (ThermoFisher Scientific) and total RNA was subsequently isolated with Direct-zol™ RNA MiniPrep (Zymo Research, Irvine, CA, USA) following the manufacturer’s instructions. Yield and purity were determined by absorbance at 230, 260, and 280 nm using NanoDrop 2000c spectrophotometer (ThermoFisher Scientific). Quantification of D2R gene expression was performed on Eco Real-Time PCR (Illumina, San Diego, CA, USA) using One Step SYBR PrimeScript RT-PCR II kit (Takara Bio, Shiga, Japan). The expression level of RP49 was used as a housekeeping (normalizing) gene. Relative gene expression was quantified with the ΔΔCt Comparative method. The primers used for expression of D2R gene were: forward primer, 5’-CCTTCTACAACGCCGACTTTA-3’, reverse primer 5’- ACTCCTCAGCGCCTTGAA-3’. To avoid eventual contamination by genomic DNA primers were designed to span an intron–exon boundary and/or RNA samples were treated with DNase.

### Human samples

The mRNA expression values are referred to DTNBP1 NM_183040 gene expression in the human postmortem dorsolateral prefrontal cortex (DLPFC) of normal subjects across lifespan. The data are available in the open access on-line application “The Brain Cloud”, which allows the query of genome-wide gene expression data and their genetic control, http://www.libd.org/braincloud. We selected the single isoforms values on the base of Illumina probes used for the quantification. The Illumina probes used to identify the human dysbindin-1 isoforms were, for Dysbindin-1A hHA - chr6:15632467-15632536, and for Dysbibdin-1C the hHC -chr6:15735609-15735678, both referred to Human assembly Mar 2006 (NCBI 36/hg18).

Caudate samples from 18 healthy control samples and 22 schizophrenia cases were obtained from the NSW Tissue Resource Center. The tissue was processed at Neuroscience Research Australia as approved by the University of New South Wales Human Research Ethics Committee (HREC 12435; Sydney, Australia). There were no significant differences found in the demographic variables of age, sex, pH, or PMI between the diagnostic groups (Supplementary Fig. S4). The rostral caudate was dissected from anatomically matched fresh frozen coronal sections cut at 60 µm through the head of the caudate. Caudate extracted samples (run in duplicates) were denatured in loading buffer 2X, and boiled for 5 min at 95°C, then the denatured samples were centrifuged at 10,000 g for 5 min. Each lane was loaded with 20 mg of total protein, as in previous studies (Talbot et al. 2011; Tang et al. 2009).

### Statistics

For animal experiments, no statistical methods were used to predetermine sample sizes, although sample sizes were consistent with those from previous studies ^20, 26, 70, 72^. No explicit randomization method was used to allocate animals to experimental groups and mice were tested and data processed by investigators blind to animal identity. Statistical analyses were performed using commercial software (STATISTICA- 13.5, StaSoft, Tulsa, OK, USA and Prism 7, GraphPad, San Diego, CA, USA). Results are expressed as mean ± standard error of the mean (SEM) throughout the manuscript. Multiple Student’s *t*-test, one-way and two-way ANOVAs were used, as appropriate. The accepted value for significance was *P* < 0.05. Newman-Keul’s test for post hoc analysis was performed, when the ANOVA highlighted a statistical significance between main effects. Data distribution was tested using the D’Agostino and Pearson normality test. Data were tested for normality before statistical analysis, by using Shapiro-Wilk, the D’Agostino and Pearson tests. The experiments reported in this work were repeated independently two to four times, using mice from at least four different generations. Numbers of mice are reported in the figure legends.

