## Supplementary Figures for "Astrocytic Regulation of Basal Ganglia Dopamine/D2-Dependent Behaviors"

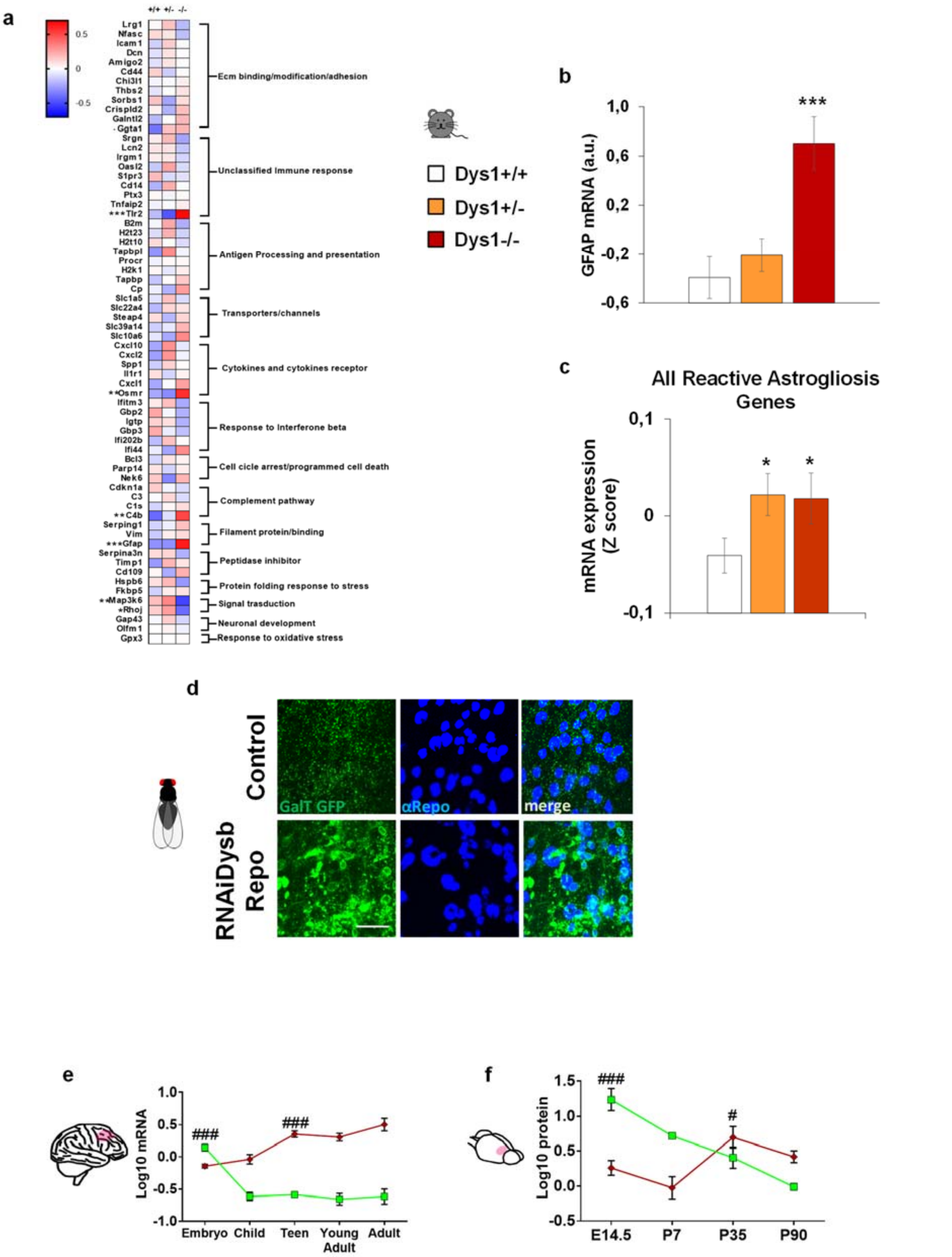


**Supplementary Figure S1. a.** Heat map of 65 inflammatory markers selected by a microarray screening from the cortex of Dys1+/+ (n24), Dys1+/- (n25), and Dys-/- (n24) littermates. The heat map is based on hierarchical clustering of genes involved in inflammation states. All gene expression levels were transformed to scores ranging from −0.5 to 0.5 and were colored blue, white, or red to represent low, moderate, or high expression levels, respectively. The relative expression levels were scaled based on their mean and do not represent expression levels in comparison with controls. **b.** Dys1-/- mice show higher GFAP expression compared to Dys1+/+ mice (One-Way ANOVA, $F_{2,66}=10.97$; $p<0.0001$). ***$p<0.0005$ vs Dys1+/+. **c.** Both Dys1+/- and Dys1-/- mice show higher expression for all the detected 65 genes involved in inflammatory process compare to Dys1+/+ littermates (One-Way ANOVA, $F_{2,190}=3.24$; $p<0.05$). *$p<0.05$ vs Dys1+/+. **d.** Maximum intensity projections of ventral ganglion cells, from *Drosophila* third instar larvae expressing UAS-GalT-GFP, of controls (repo-Gal4/+) and UAS-Dysb RNAi, to visualize Golgi cisternae in glial cells. Tissues were labeled with anti αRepo antibody to visualize glial nuclei. Scale bar 20 μm. **e.** mRNA expression of Dys1A and Dys1C isoforms from the human dorsolateral PFC by the open-access Brain Cloud databank at different developmental ages. Ns: Embryos=38; Child=32; Teen=50; Young adult=25; Adult=122. Dys1A expression was highest at the embryonic stage and then decreased, while Dys1C expression increased from adolescence (Two-way ANOVA, isoforms*age interaction: $F_{5,250}=68.84$; $p<0.0001$). $^{###}p<0.0001$ vs consecutive ages for Dys1A, and vs preceding ages for Dys1C. **f.** Protein expression of Dys1A and Dys1C isoforms from the mouse prefrontal cortex at different developmental ages. n6 mice/time point. Dys1A expression was highest at the embryonic stage and then decreased, while Dys1C increased its expression from adolescence (Two-way ANOVA, Isoforms*Age interaction: $F_{3,17}=32.77$; $p<0.0001$). $^{\#}p<0.01$ and $^{###}p<0.0001$ vs consecutive ages for Dys1A, and vs preceding ages for Dys1C. Bar graphs show mean ± s.e.m.

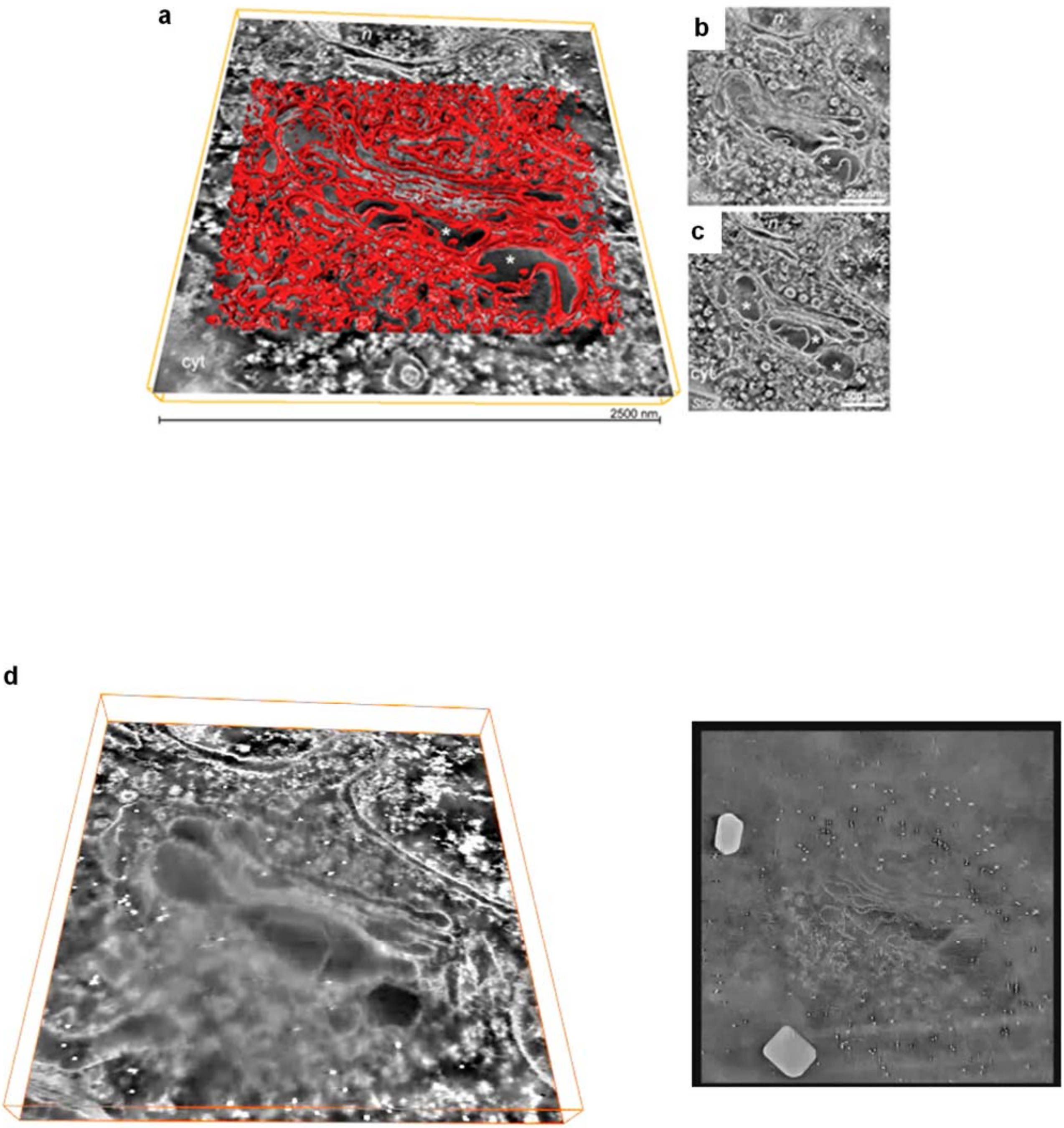


**Supplementary Video V1.** Scanning TEM (STEM) dark field double tilt electron tomography of an altered Golgi complex of a Dys1+/- mouse astrocyte. **a.** Tomogram 3D reconstruction (in red) superimposed to a single tomographic slice. **b-c.** Different slices through the tomogram. The asterisks point to swollen and irregularly shaped Golgi cisternae. Cyt: cytoplasm; n: nucleus. **d.** Series of slices through a reconstructed dark field STEM tomogram of an altered Golgi complex of a Dys1+/- mouse astrocyte with its 3D reconstruction. The side of the tomogram field of view is equal to 2.6 um.

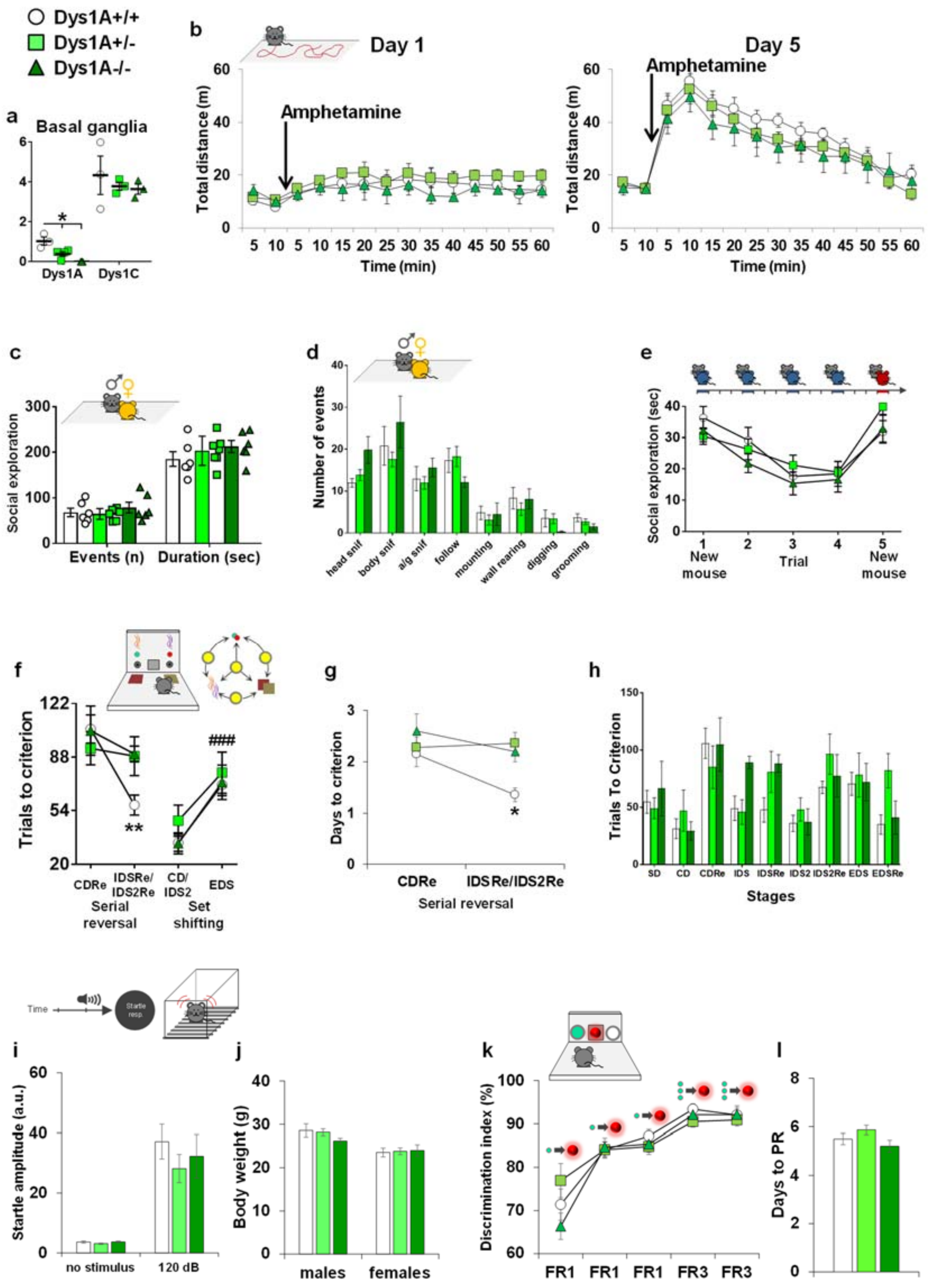


**Supplementary Figure S2. a.** Selective reduction of Dys1A isoform does not affect Dys1C expression. Western blots and densitometric analysis in Dys1A+/+, +/- and -/- littermates. Expression of Dys1A (50 kDa), Dys1C (38 kDa), and β-actin (42 kDa) as loading control. Expression of Dys1A is reduced in Dys1A+/- and absent in Dys1A-/- in the basal ganglia regions (One-way ANOVA, $F_{2,7}$=14.29; p<0.005). *p<0.01 vs Dys1A+/+ littermates. Expression of Dys1C was intact across all genotypes (One-way ANOVA, $F_{2,7}$=0.41; p=0.68). n3-4 mice/group. **b.** Dys1A knockout mice unaltered sensitization to amphetamine (a) Acute exposure to amphetamine and sensitization to amphetamine following five days of administration displayed by Dys1A+/+ (n15), Dys1A+/- (n16) and Dys1A-/- (n10) exposed to the open field apparatus. All the groups were equally sensitized to amphetamine injection independently by the genotype (day1 vs day 5 p<0,0005). **c.** Male-female social interaction, displayed by Dys1A+/+ (n6), Dys1A+/- (n7) and Dys1A-/- (n6) littermates. No genotype-dependent difference was evident in the number of events or exploration time (Two-way ANOVA; events: $F_{2,15}$=0.52; p=0.60; duration: $F_{2,15}$=0.75; p=0.49). **d.** Male/female social interaction. Frequency of occurrence of various social behaviors such as head sniffing, body sniffing, anogenital sniffing and following, and non-social events including standing/walking alone, digging, grooming, rearing and wall rearing displayed by Dya1A+/+ (n6), Dys1A+/- (n7) and Dys1A-/- (n6) littermates. No significant genotype effect was evident in any parameter measured. **e.** Social habituation-dishabituation, displayed by Dys1A+/+ (n14), Dys1A+/- (n16) and Dys1A-/- (n10) littermates. No genotype-dependent difference was evident (Two-way repeated measure ANOVA genotype: $F_{2,37}$=0.37; p=0.69; genotype*time interaction: $F_{8,148}$=1.88; p=0.07). **f.** Attentional Set-Shifting Test (ASST), performed in Dys1A+/+ (n14), Dys1A+/- (n14) and Dys1A-/- (n10) littermates. In contrast to Dys1A+/+, both Dys1A+/- and Dys1A-/- mice did not show the expected reduced trials to criterion in serial reversal stages (Two-way repeated measure ANOVA; reversal stage: $F_{1,35}$=11.46; p=0.002; genotype*reversal stage interaction: $F_{2,35}$=4.27; p=0.02). **p<0.005 vs Dys1A+/- and -/- at the same time point. All mice showed the expected increase in trials to criterion at EDS stage independent of genotype (Two-way repeated measures ANOVA; shifting stage: $F_{1,35}$=17.45; p=0.0002; genotype*shifting stage interaction: $F_{2,35}$=0.09; p=0.92). ###p<0.0005 vs CD/IDS2 stages. **g.** Days needed to reach criterion in ASST stages shown by Dys1A+/+ (n14), Dys1A+/- (n14) and Dys1A-/- (n10) littermates. In contrast to Dys1A+/+ mice, both Dys1A+/- and Dys1A-/- mice did not show the expected reduced days to criterion in serial reversal stages (Two-way repeated measure ANOVA; reversal stage: $F_{1,35}$=5.01; p=0.03; genotype*reversal stage interaction: $F_{2,35}$=2.60; p=0.05). **h.** Total amount of trials needed to react the criteria for each stage of the ASST displayed by Dys1A+/+ (n14), Dys1A+/- (n14) and Dys1A-/- (n10) littermates. **i.** Analysis of 120 dB acoustic startle stimulus showed no interaction between genotype Dys1A+/+ (n10), Dys1A+/- (n15) and Dys1A-/- (n12) and startle response ($F_{2,34}$=0,495; p=0,613) In all the groups we found a main effect of startle response vs no stimulus ($F_{1,34}$=71,42; p<0,0001). **j.** Body weight displayed by the same Dys1A+/+ (n10), Dys1A+/- (n15) and Dys1A-/- (n12) performing the PPI test divided by sex. **k.** Last five days of training in the fixed ratio 1 and 3 schedule before starting the progressive ratio test displayed by Dys1A+/+ (n12), Dys1A+/- (n13) and Dys1A-/- (n10) littermates. **l.** Days needed to reach the final progressive ratio test (PR) stage displayed by the same Dys1A+/+ (n12), Dys1A+/- (n13) and Dys1A-/- (n10) littermates. Bar and line graphs show mean ± s.e.m.

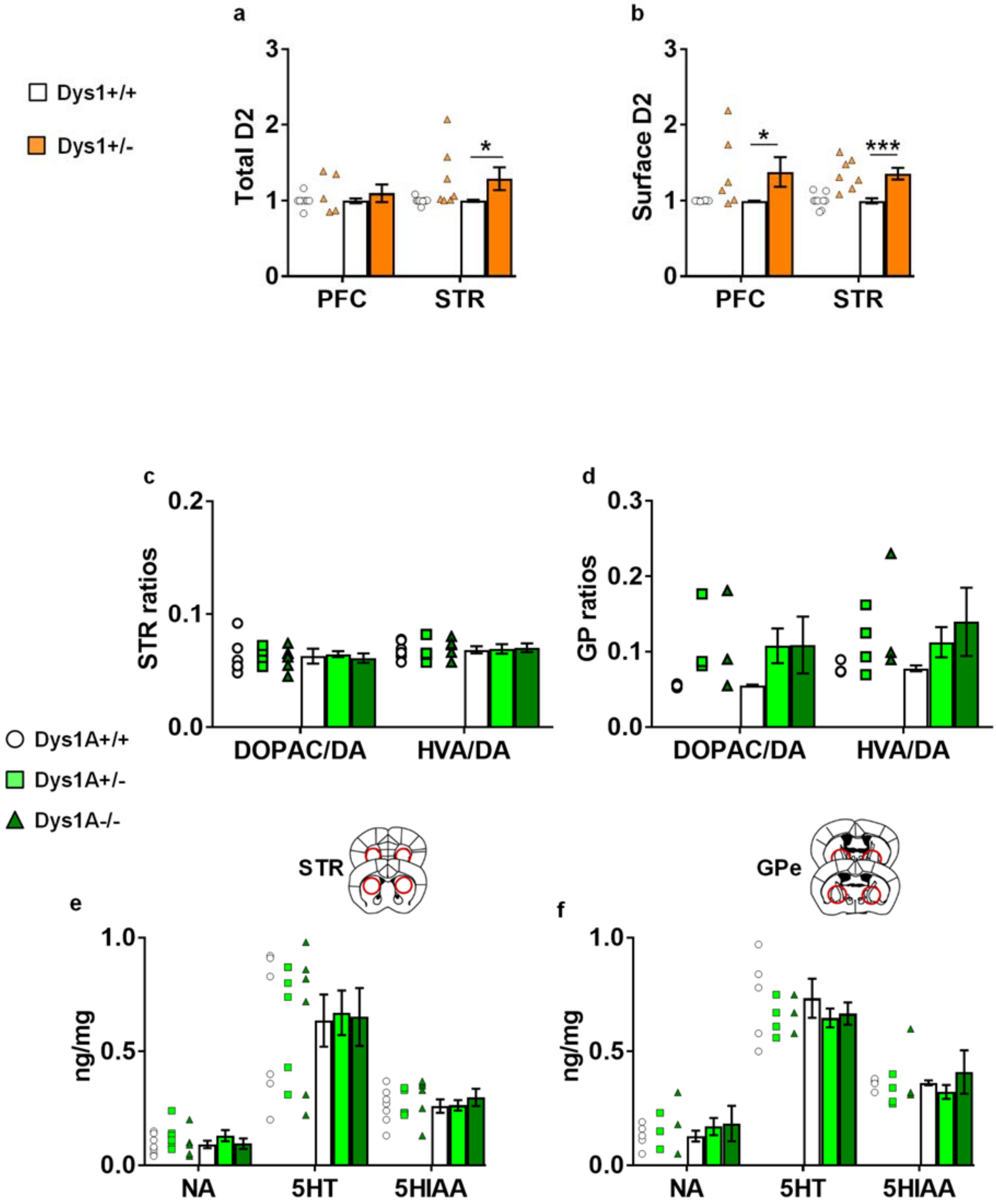


**Supplementary Figure S3. a.** D2 receptor expression in Dys1+/+ (n9), and Dys1+/- (n7), synaptophysin has been used as cytoplasmic control, D2 expression has been normalized on transferrin as loading control. Dys1+/- mice had increased total D2 expression compared to Dys1+/+ in the STR (t-test: $t_{14}$=-2.19, p=0.046), but not in the PFC (t-test: $t_{14}$=-2.19, p=0.046). *p<0.05 vs Dys1+/+. **b.** Biotinylation of brain slices treated with biotin to label all surface proteins, precipitated by streptavidin. Dys1+/- mice had increased expression of D2 receptors on the cellular surface compared to Dys1+/+ littermates in the PFC and STR (PFC: t-test: $t_{12}$=-2.27, p=0.042; STR: t-test: $t_{14}$=-4.65, p=0.0003). ***p<0.0005 and *p<0.05 vs Dys1+/+. **c-d.** DOPAC/dopamine and

HVA/dopamine ratios from HPLC experiments in the **c.** STR, and **d.** GPe dissected from Dys1A+/+ (n5), Dys1A+/- (n4), and Dys1A-/- (n5) littermates. No Dys1A-dependent changes were observed (One-way ANOVAs, STR DOPAC/DA: $F_{2,15}$=0.14; p=0.87; STR HVA/DA: $F_{2,14}$=0.05; p=0.95; GPe DOPAC/DA: $F_{2,8}$=1.91; p=0.21; GP HVA/DA: $F_{2,8}$=1.54; p=0.27). **e-f.** Noradrenaline (NA), serotonin (5HT), and 5-hydroxyindoleacetic acid (5HIAA) content by HPLC analyses, expressed as ng/mg of tissue in the **e.** STR, and **f.** GPe dissected from Dys1A+/+ (n6), Dys1A+/- (n5), and Dys1A-/- (n6) littermates. No Dys1A-dependent changes were observed (One-way ANOVAs, STR NA: $F_{2,16}$=1.00; p=0.39; STR 5HT: $F_{2,16}$=0.02; p=0.98; STR 5HIAA: $F_{2,16}$=0.39; p=0.68; GPe NA: $F_{2,9}$=0.48; p=0.63; GPe 5HT: $F_{2,9}$=0.46; p=0.65; GPe 5HIAA: $F_{2,9}$=0.87; p=0.45). Bar and line graphs show mean ± s.e.m.

| | Patients with Schizofrenia | Healthy Control | t-value | df | p |
|---|---|---|---|---|---|
| **N° of Cases** | 22 | 18 | | | |
| **Male/Female** | 13/9 | 15/3 | | 1 | 0,096 |
| **Mean Age ± SD** | 55,18±13,54 | 55,21±14,1 | -0,006 | 39 | 0,997 |
| **Mean tissues pH ± SD** | 6,6±0,27 | 6,64±0,23 | -0,505 | 39 | 0,615 |
| **Mean PMI ± SD** | 32,75±14,02 | 29,81 ± 10,06 | 0,758 | 39 | 0,452 |

**Supplementary Figure S4.** Demographic and autopsy data on postmortem cases studied.

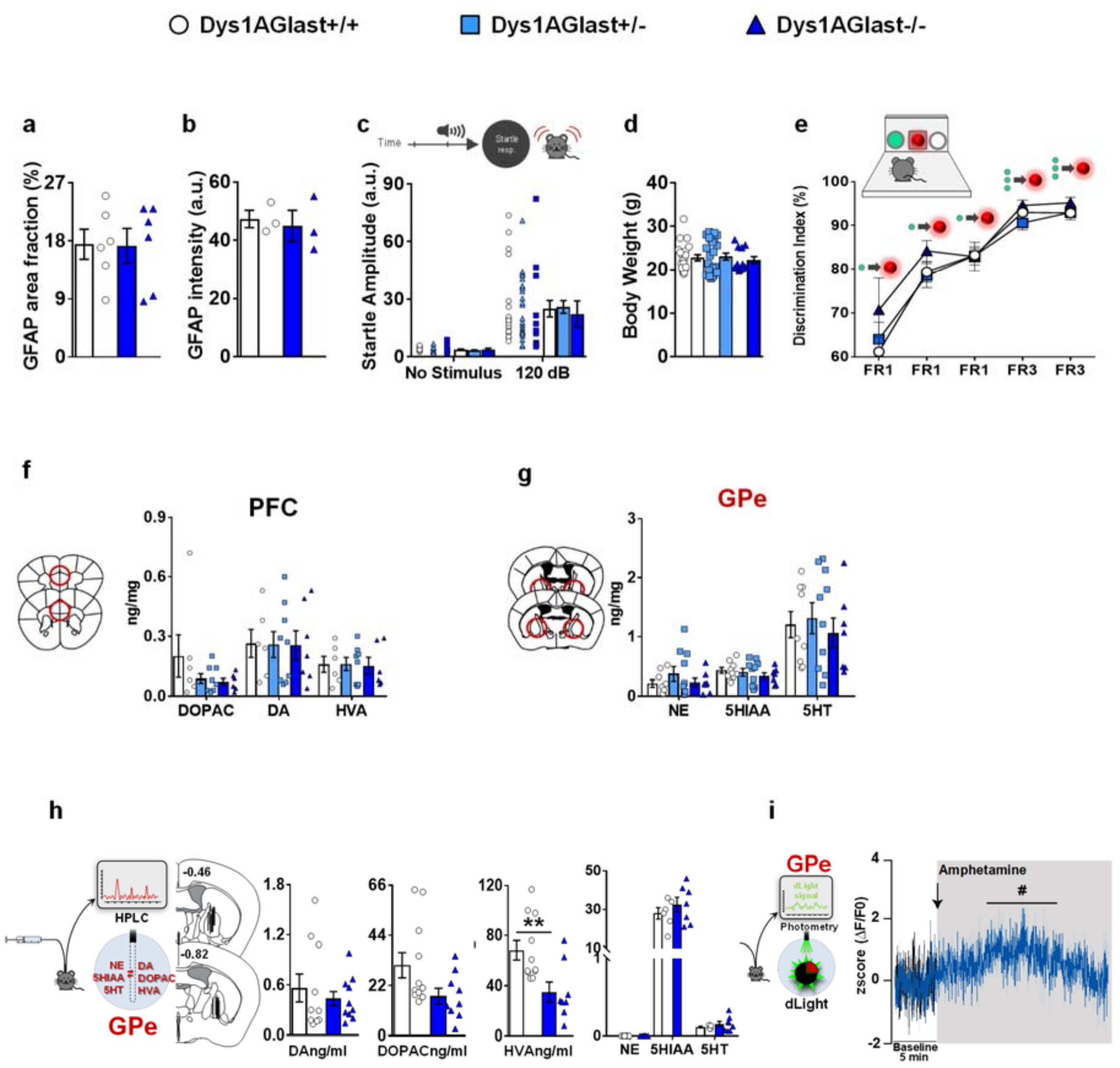


**Supplementary Figure S5. a.** There was no difference in GFAP stained area between genotypes (t-test: $t_{10}$=0.06, p=0.95). **b.** There was no difference in GFAP intensity between genotypes (t-test: $t_4$=0.39, p=0.72). **c.** Analysis of 120 dB acoustic startle stimulus showed no interaction between genotype Dys1AGlast+/+ (n21), Dys1AGlast+/- (n24) and Dys1AGlast-/- (n12) and startle response ($F_{2,54}$=0,20; p=0,82). In all groups we found a main effect of startle response vs no stimulus ($F_{1,54}$=64,12; p<0,0001). **d.** Body weight displayed by the same Dys1AGlast+/+ (n21), Dys1AGlast+/- (n24) and Dys1AGlast-/- (n12) performing the PPI test (genotype effect: $F_{2,54}$=0.22; p=0.80). **e.** Last five days of training in the fixed ratio 1 and 3 schedule before starting the progressive ratio test displayed by Dys1AGlast+/+ (n16), Dys1AGlast+/- (n16) and Dys1AGlast-/- (n10) littermates. All mice improved their performance in a genotype-independent way (genotype effect: $F_{2,39}$=0.53; p=0.60; day effect: $F_{4,156}$=88.87; p<0.0001; genotype-day interaction: $F_{8,156}$=1.34; p=0.23). **f.** Dopamine (DA), DOPAC, and HVA content by HPLC analyses, expressed as ng/mg of dissected PFC tissues displayed by Dys1AGlast+/+ (n6), Dys1AGlast+/- (n9), and Dys1AGlast-/-

(n7) littermates. No genotype-dependent effects were evident (One-way ANOVAs, DOPAC: $F_{2,17}$=1.35; p=0.29; DA: $F_{2,19}$=0.21; p=0.81; HVA: $F_{2,17}$=0.02; p=0.98). **g.** Norepinephrine (NE), 5-Hydroxyindoleacetic acid (5HIAA), and serotonin (5HT) content by HPLC analyses, expressed as ng/mg of dissected GPe tissues displayed by Dys1AGlast+/+ (n9), Dys1AGlast+/- (n10), and Dys1AGlast-/- (n7) littermates. No genotype-dependent effects were evident (One-way ANOVAs, NE: $F_{2,23}$=0.95; p=0.40; 5HIAA: $F_{2,23}$=0.60; p=0.56; 5HT: $F_{2,23}$=0.24; p=0.79). **h.** No genotype-dependent differences in basal extracellular dopamine levels (t-test: $t_{18}$=0.67, p=0.51), and DOPAC levels (t-test: $t_{18}$=0.32, p=0.75). Decreased extracellular HVA levels in GPe of Dys1AGlast-/- mice compared to Dys1AGlast+/+ littermates (t-test: $t_{18}$=2.95, p=0.009). **p<0.005 vs Dys1AGlast+/+. Basal extracellular norepinephrine (NE), 5-Hydroxyindoleacetic acid (5HIAA), and serotonin (5HT) content by HPLC analyses, expressed as ng on ml from microdialysis probes for measurement of extracellular levels displayed by Dys1AGlast+/+ (n6), and Dys1AGlast-/- (n8) littermates. No genotype-dependent effects were evident (One-way ANOVAs, NE: undetectable in all samples; 5HIAA: $F_{1,11}$=0.88; p=0.37; 5HT: $F_{1,11}$=0.85; p=0.38). **i.** Analyses of the dLight signal in the GPe. Amphetamine (1.5mg/kg, i.p.) equally increased dLight signal compared to the 5-min baseline period before injection in the GPe of both Dys1AGlast+/+ and Dys1AGlast-/- littermates (Two-way RM ANOVA; genotype: $F_{1,4}$=1.35; p=0.31; Time: $F_{19,76}$=1.86; p<0.05). #p<0.05 vs their own baseline.

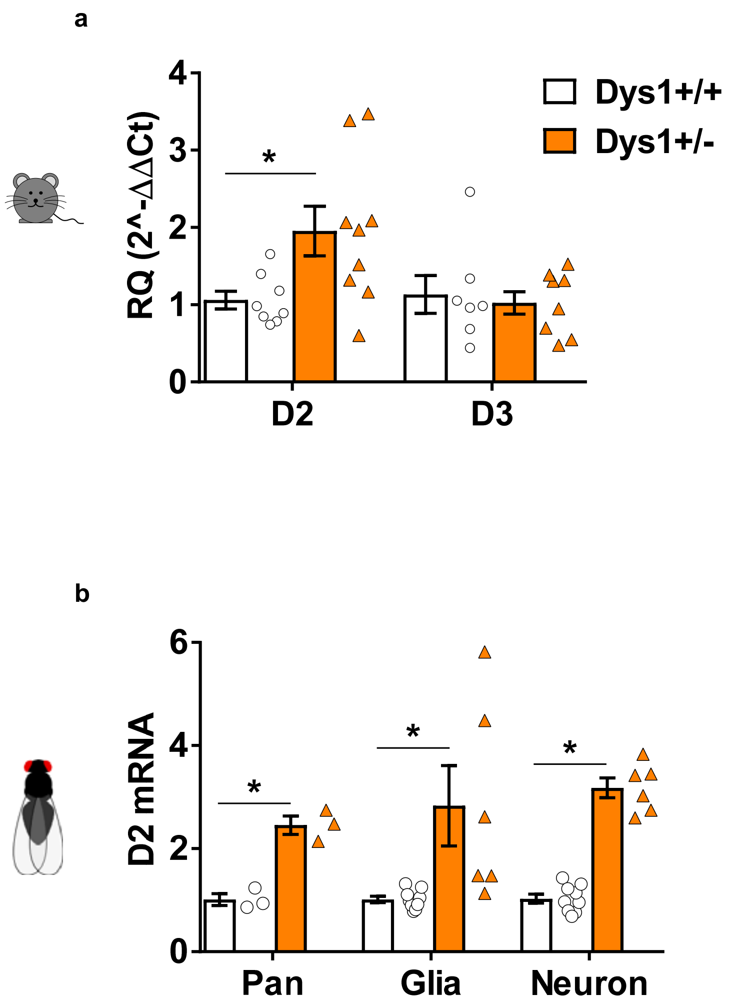


**Supplementary Figure S6. a.** Abundance of dopamine D2 and D3 receptors in the STR + GPe regions displayed by Dys1+/+ (n8) and Dys1+/- (n9) measured by quantitative RT-PCR. t-test *p<0.05 vs Dys1+/+. **b.** Drosophila third instar larvae, control Pan (tubulin-Gal4/+), glial (Repo-Gal4/+); neuronal (Elav-Gal4/+) and RNAi Dysb/tubulin-Gal4; glial RNAi Dysb/Repo-Gal4/+; neuronal (Elav-Gal4/+). Relative mRNA expression of D2 receptors. t-test *p<0.05 vs control group. Bar and line graphs show mean ± s.e.m.

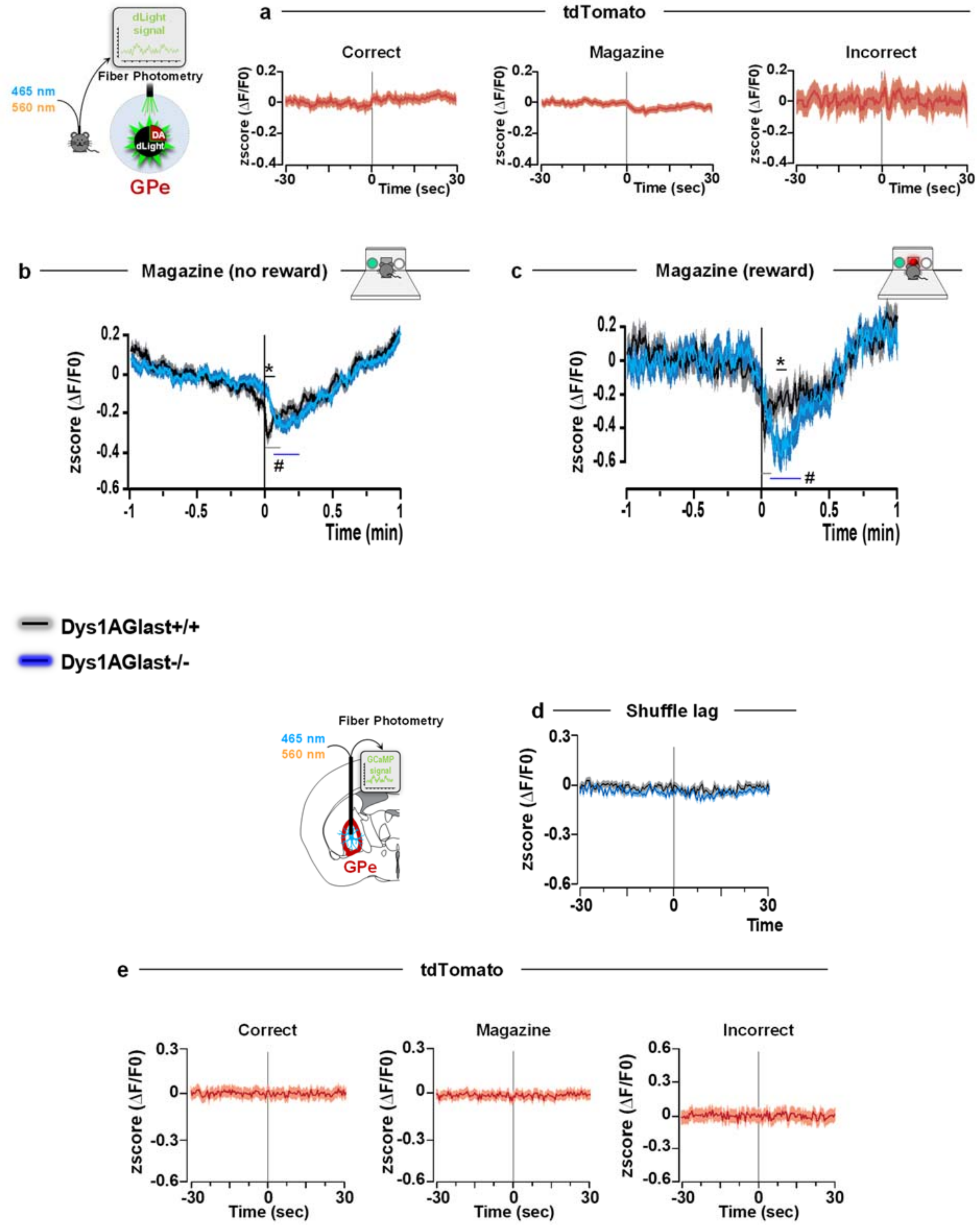


**Supplementary Figure S7. a.** Analyses of the tdTomato signal in the GPe during the correct, magazine, and incorrect actions revealed no ΔF/F0 changes compared to baseline values in the dLight fiber photometry experiment. **b.** Analyses of the dLight signal in the GPe during food magazine entrance in absence of pellet delivery (Magazine, no reward) revealed a decreased (ΔF/F0) signal in Dys1AGlast+/+ from the entrance to the magazine up to 7 sec later, and from 3 sec up to 17 in Dys1AGlast-/- mice. A genotype-dependent difference between Dys1AGlast+/+ and Dys1AGlast-/-

mice was evident from magazine entrance to 2.5 sec later. #p<0.05 vs own genotype baseline. *p<0.05 vs Dys1AGlast+/+ at the same time interval. **c.** Analyses of the dLight signal in the GPe during food magazine entrance in presence of food pellet (Magazine, reward) revealed a decreased (ΔF/F0) signal in Dys1AGlast+/+ from the entrance to the magazine up to 1.5 sec later, and from 2.5 sec up to 16.5 in Dys1AGlast-/- mice. A genotype-dependent difference between Dys1AGlast+/+ and Dys1AGlast-/- mice was evident from 8 to 9 sec after magazine entrance. #p<0.05 vs own genotype baseline. *p<0.05 vs Dys1AGlast+/+ at the same time interval. **d.** Astrocytes $Ca^{2+}$ signal responses by fiber photometry in GPe of Dys1AGlast+/+ and Dys1AGlast-/- mice synchronized at shuffle times revealed no effects. **e.** Analyses of the tdTomato (tdT) control signal in GPe during correct, magazine, and incorrect actions revealed no ΔF/F0 changes compared to baseline values. Bar and line graphs show mean ± s.e.m.

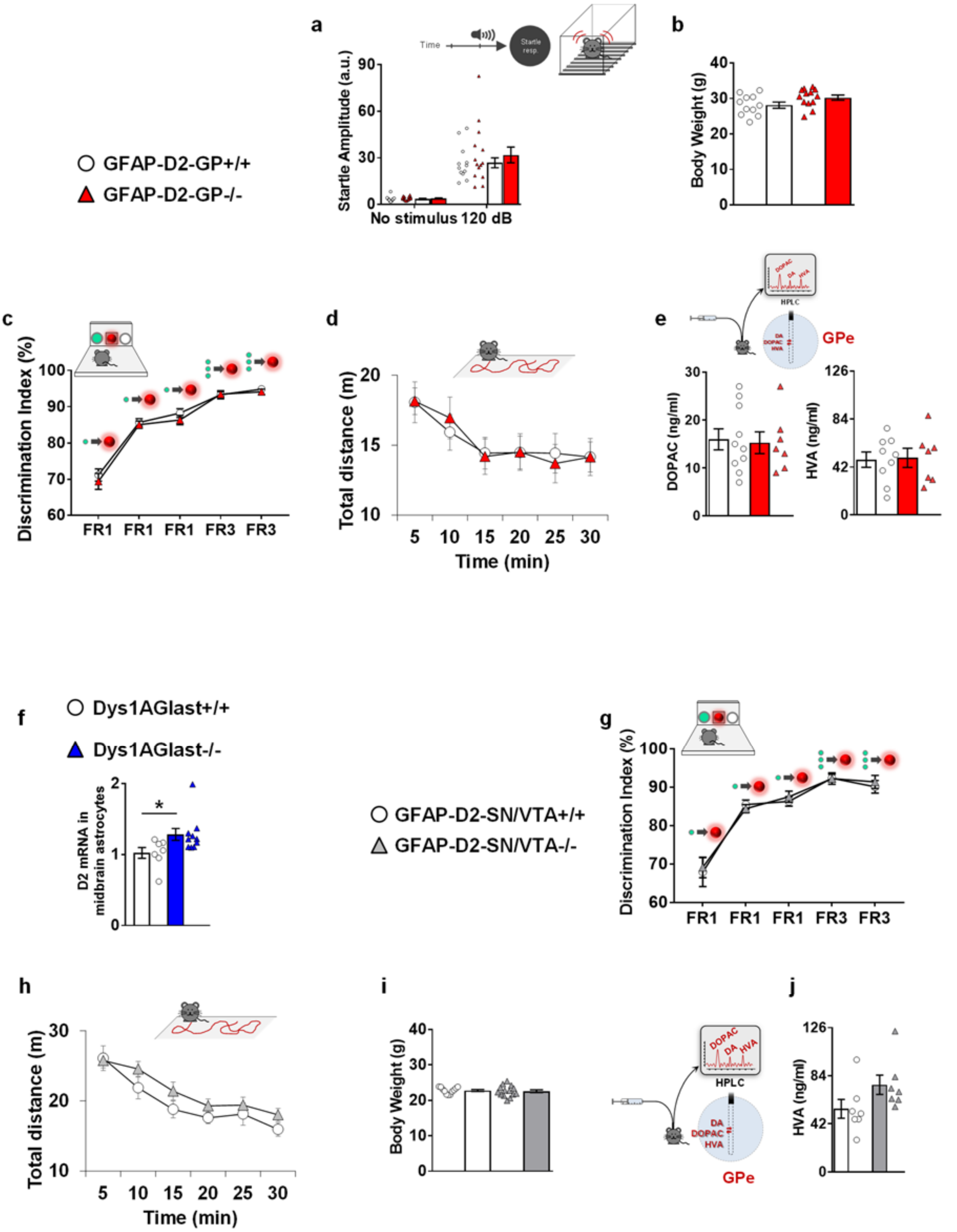


**Supplementary Figure S8. a.** Analysis of 120 dB acoustic startle stimulus showed no interaction between genotype (GFAP-D2-GP+/+, n12; GFAP-D2-GP-/-, n14) and startle response ($F_{2,54}$=0,20; p=0,82). In all groups we found a main effect of startle response vs no stimulus ($F_{1,54}$=64,12; p<0,0001). **b.** Body weight displayed by the same GFAP-D2-GP+/+ (n12) and GFAP-D2-GP-/- (n14) performing the PPI test (genotype effect: $F_{1,22}$=3.60; p=0.08). **c.** Last five days of training in the fixed ratio 1 and 3 schedule before starting the progressive ratio test displayed by GFAP-D2-GP+/+ (n12)

and GFAP-D2-GP-/- (n12) littermates. All mice improved their performance in a genotype-independent way (genotype effect: $F_{1,21}$=1.19; p=0.29; day effect: $F_{4,84}$=122.18; p<0.0001; genotype-day interaction: $F_{4,84}$=0.17; p=0.95). **d.** Spontaneous distance traveled by GFAP-D2-GP+/+ (n14) and GFAP-D2-GP-/- (n16) during 30 min exposure to an open field arena. No genotype differences were evident (Two-way repeated measure ANOVA, genotype effect: $F_{1,28}$=0.00; p=0.99; time*genotype interaction: $F_{5,140}$=0.40; p=0.85). **e.** GFAP-D2-GP+/+ (n10) and GFAP-D2-GP-/- (n7) littermates were implanted with a dialysis probe for measurement of basal extracellular DOPAC and HVA levels within the GPe. No dependent differences were evident between groups (t-test; DOPAC: t=0.55, df=16, *p*=0.59; HVA: t=0.66, df=16, *p*=0.52). **f.** Relative mRNA expression of dopamine D2 receptors, assessed by RT-qPCR in tdTomato-positive cells sorted from the midbrain of Dys1AGlast+/+ (n7) and Dys1AGlast-/- (n10) littermates, revealing an increased astrocytic D2 receptor expression in DysGlast-/- mice (t-tests: $t_{15}$=-2.21, p=0.043). Data shown as fold-change compared with Dys1AGlast+/+ control mice. *p<0.05 vs Dys1AGlast+/+. **g.** Last five days of training in the fixed ratio 1 and 3 schedule before starting the progressive ratio test displayed by GFAP-D2-SN/VTA+/+ (n12) and GFAP-D2-SN/VTA-/- (n14) littermates. All mice improved their performance in a genotype-independent way (genotype effect: $F_{1,24}$=0.09; p=0.77; day effect: $F_{4,96}$=70.38; p<0.0001; genotype-day interaction: $F_{4,96}$=0.21; p=0.93). **h.** Spontaneous distance traveled by GFAP-D2-SN/VTA+/+ (n10) and GFAP-D2-SN/VTA-/- (n14) during 30 minutes exposure to an open field arena. No genotype differences were evident (Two-way repeated measure ANOVA, genotype effect: $F_{1,22}$=0.99; p=0.33; time*genotype effect: $F_{5,110}$=1.48; p=0.20). **i.** Body weight displayed by GFAP-D2-SN/VTA+/+ (n11) and GFAP-D2-SN/VTA-/- (n14) littermates (t=0.40, df=23, *p*=0.69). **j.** GFAP-D2-SN/VTA+/+ (n7) and GFAP-D2-SN/VTA-/- (n7) littermates were implanted with a dialysis probe for measurement of basal extracellular dopamine, DOPAC and HVA levels within the GPe. No genotype-dependent differences were evident for HVA (t-test: t=-1.78, df=12, *p*=0.10). Bar and line graphs show mean ± s.e.m.
